# Region-specific spreading depolarization drives aberrant post-ictal behavior

**DOI:** 10.1101/2024.10.12.618012

**Authors:** Bence Mitlasóczki, Adrián Gutiérrez Gómez, Midia Kamali, Natalia Babushkina, Mayan Baues, Laura Kück, André Nathan Haubrich, Theodoros Tamiolakis, Annika Breuer, Simon Granak, Merlin Schwering-Sohnrey, Ingo Gerhauser, Wolfgang Baumgärtner, Laura Ewell, Thoralf Opitz, Julika Pitsch, Simon Musall, Rainer Surges, Florian Mormann, Heinz Beck, Michael Wenzel

## Abstract

Confusion, aphasia, and unaware wandering are prominent post-ictal symptoms regularly observed in temporal lobe epilepsy (TLE)^1^. Despite the potentially life-threatening nature of the immediate post-ictal state^2^, its neurobiological underpinnings remain understudied^3^. We provide evidence in mice and humans that seizure-associated focal spreading depolarization (sSD) is a pathoclinical key factor in epilepsy. Using two-photon or widefield imaging (hippocampus, neocortex), field potential and single unit recordings, and behavioral assessment in mice, we first studied seizures during viral encephalitis, and subsequently established an optogenetic approach to dissociate hippocampal seizures and SD. We find region-specific occurrence of sSD that displays distinct spatial trajectories to preceding seizures, and show that seizure-related and isolated hippocampal SD prompt *post-ictal wandering*. This clinically relevant locomotor phenotype occurred in the absence of hippocampal SD progression to the neocortex. Finally, we confirm sSD existence in human epilepsy, in a patient cohort with refractory focal epilepsy, via Behnke-Fried electrode recordings. In this cohort, sSD displayed a similar temporomesial propensity as in mice. This work uncovers sSD as a previously underrecognized pathoclinical entity underlying postictal behavioral abnormalities in epilepsy. Our results carry wide-reaching ramifications for epilepsy research and neurology, and challenge current EEG-standards.

## Introduction

The immediate post-ictal period often manifests with confusion, aphasia, amnesia, or unaware ambulation (*post-ictal wandering*), which is most commonly observed in temporal lobe epilepsy (TLE)^1,3^. It is associated with a risk for severe, potentially life-threatening injuries^2^. Despite its clinical impact, except for particular entities such as sudden unexpected death in epilepsy (SUDEP)^4,5^, the neurobiological underpinnings of the immediate post-ictal state remain unclear, and most related research on this topic has focused on consequence rather than initial cause^6,7^. One reason for the continued scarcity of neurobiological insight may be that the *ictus* itself has continually formed the center of interest in epilepsy research and clinical care, which is also reflected in clinical terminology (ictus, ictal, pre-/post-/peri-ictal).

Previous work has suggested that network dynamics other than seizures (Sz), e.g. spreading depolarization/depression (SD)^8^, could in principle account for a number of post-ictal symptoms^3,4,9^. Most of what is known about SD stems from research on migraine, stroke and traumatic brain injury^10–12^. In the latter context, Sz and neocortical SD have been investigated also in clinical electrical recordings^13–22^. While generally, SD has been recorded in hippocampus^23–29^, thalamus^30,31^, basal ganglia^23^, or brainstem^4,5^, the vast majority of SD research has focused on neocortex, which brought about the common term cortical spreading depression (CSD)^32^. Put simply, SD constitutes a massive ion translocation across the neuronal cell membrane resulting in a profound depolarization above the inactivation threshold, a break-down of the membrane potential and consecutive neuronal depression^8,32,33^. SD can be elicited in various ways, e.g. by energy depletion^34–36^, hypoxia^37^, high extracellular K^+36,38–40^, repetitive electrical or mechanical stimulation^8^, or optogenetic neuronal depolarization^41–46^. Depending on the etiology, SD has profound effects on affected brain tissue such as transient edema and vasoconstriction, danger molecule release in the extracellular space (i.e. ATP), or immune activation^12,26,37,47^. Such effects may be transient and recovered from, but can also lead to permanent tissue damage^21,48^.

While some of the mentioned studies have documented an association of SD and Sz in acute brain injury in neurointensive care^15,18^, it is unknown whether Sz-related SD exists in human epilepsy at the clinical level. In basic epilepsy research, the role of SD remains debated. SD has both been suggested to increase neural excitability^29,49^, and to disrupt ictal oscillations^27,50,51^. Some have speculated about a protective role of SD in epilepsy^50,52^, while others have described Sz-related SD as a potential cause for SUDEP via brain stem invasion^4,5^. Certainly, as most basic research on Sz-related SD has been carried out in vitro, or under anesthesia^8,9,23,26,49,50^, the general relationship between Sz-related SD and potentially distinct clinical symptoms have remained understudied. In current clinical epileptology, concepts revolving around Sz-related SD, aside from SUDEP, play little to no role.

Here, we provide *in-vivo* evidence in mice and humans that Sz-associated focal spreading depolarization (sSD) constitutes a proper pathoclinical entity in epilepsy. Using two-photon or one-photon widefield imaging (hippocampus, neocortex), field potential and single unit recordings, and behavioral assessment in mice, we first studied seizures during viral encephalitis, and subsequently established an optogenetic approach to dissociate hippocampal seizures and SD. We find profound region-specific differences in sSD occurrence, characterize its progression, and show that temporomesial SD triggers *post-ictal wandering*. Then, by Behnke-Fried (BF) electrode recordings in an initial cohort of four patients with refractory focal epilepsy undergoing pre-surgical diagnostic work-up, we provide corroborating evidence that region-specific sSD exists in human focal epilepsy too, with a strikingly similar regional propensity distribution of occurrence as observed in our murine models.

## Results

### Two-photon Ca^2+^ imaging of epileptic network dynamics during viral encephalitis

We initially set out to study naturally occurring temporal lobe Sz at cellular scale. To this end, we employed resonant two-photon (2p) population Ca^2+^ imaging in the hippocampus (CA1) or cortex (CTX, motor) of awake adult transgenic Thy1-GCaMP6s mice (JAX 025776; 15 Hz scanning, ∼700x700 µm, 16X Nikon 0.80 NA 3.0 mm WD) (Fig. 1A, B), in the recent Theiler’s murine encephalomyelitis virus (TMEV) etiocopy mouse model of TLE. The model is based on an initial self-limiting viral encephalitis that encompasses acute disease stage focal onset seizures (∼2-7 days post infection [p.i.], Fig. 1 C, Suppl. Fig. 1, for model details, see methods)^53^.

**Fig. 1.**
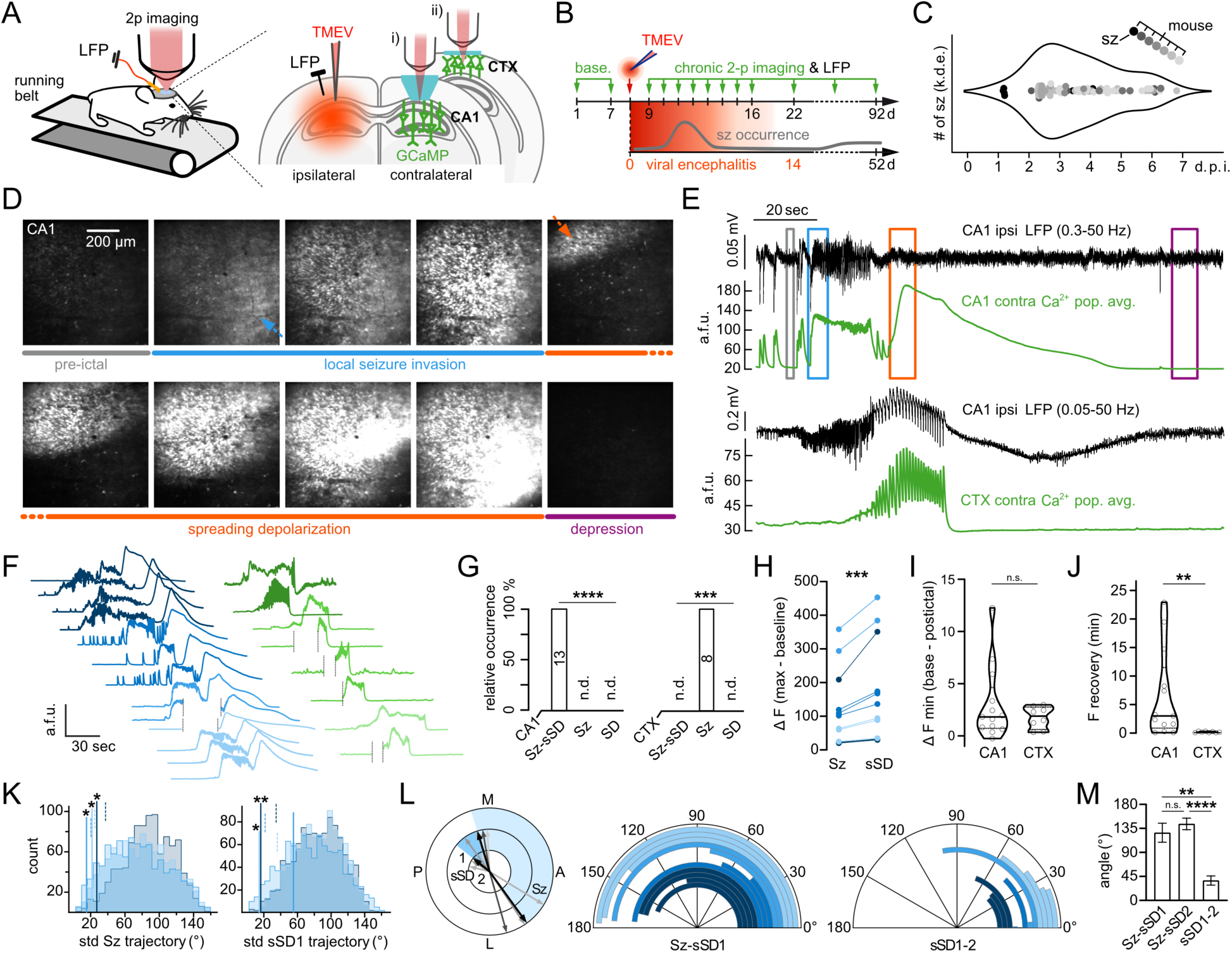
Chronic hippocampal or neocortical 2p-imaging of seizures during viral encephalitis. **A**, Exp. setup. Contralateral (contra) cranial window above hippocampus (i: CA1) or neocortex (ii: CTX) to ipsilateral (ipsi) LFP electrode (black pin, at CA1) and TMEV injection site (CA1), in transgenic GCaMP6s mice. **B**, Exp. workflow. **C**, Detected clinical seizures (Sz, filled circles) during viral encephalitis in initial video-monitoring (7 mice, gray shades). Max. number (#) of Sz (k.d.e.: kernel density estimate) typically occur on day 2-5 post injection (d.p.i.). **D**, Representative average (avg) fluorescence (F) images of neuronal signals during pre-ictal baseline (gray), Sz (blue) and sSD (red/violet) invasion of CA1. Dotted arrows depict travelling direction of Sz (blue) or sSD (red). **E**, LFP (black, ipsi CA1) and avg population Ca^2+^ signals (green, contra) in CA1 (top) or CTX (bottom) during local Sz and sSD invasion (a.f.u.: arbitrary fluorescent units). Colored boxes correspond to time periods for avg images in D. Note that sSD is not detected in LFP with a 0.3 Hz high-pass (HP) filter (top panel), whereas a large biphasic shift is visible in bottom panel (0.05 Hz HP). Note also delayed optical CTX Sz invasion. **F**, All imaged Sz in CA1 (13 Sz, 4 mice, blue shades) or CTX (8 Sz, 3 mice, green shades). Note the presence or absence of sSD. Dotted lines indicate momentary breaks between imaging sessions. **G**, Quantification of occurrence of Sz-sSD, Sz, or SD during encephalitis in CA1 (left, Friedman test, p<0.0001) or CTX (right, Friedman test, p=0.0005), n.d. = not detected. **H**, Comparison of Sz vs SD relative Ca^2+^ signal amplitude in CA1 (a.f.u., ∆ F: Delta Fluorescence), paired Wilcoxon test (12 Sz-sSD, 117.1 ± 32.6 [Sz] vs. 164.8 ± 43.35 [sSD], p=0.0005). **I**, Comparison of CA1 vs CTX pre-and post-ictal minimal F (a.f.u., ∆ F min), Mann-Whitney test (13 events in CA1 vs. 8 events in CTX, 3.02 ± 3.54 vs. 1.77 ± 1.04, p=0.8044). **J**, Comparison of CA1 vs CTX F recovery time (minutes), Mann-Whitney test (13 events in CA1 vs. 8 events in CTX, 6.531 ± 2.15 vs. 0.188 ± 0.038, p=0.0061). **K**, Observed (solid lines) standard deviations (std) of spatial trajectory angles (°) of Sz (left) or sSD1 (right) from 3 mice (blue shades, each mouse z 3 imaged Sz-sSD) vs corresponding shuffled distributions from 1000 randomized datasets. Dashed lines mark respective significance threshold (observed std <95% of all surrogate std). **L**, Left: Paradigmatic experiment with spatial Sz-sSD trajectory map within imaged field of view (M medial, L lateral, P posterior, A anterior). Outer ring shows Sz, intermediate ring sSD1, inner ring sSD2 trajectories (gray shades: individual events, black: mean). Shaded areas display trajectory angle between Sz and sSD1 (light blue), or sSD1 and sSD2 (dark blue). Middle and Right: Calculated angles (°) for each imaged Sz-sSD1 and sSD1-sSD2. One event excluded from analysis, as immediate Sz onset was not recorded, see Fig. 1 F. **M**, Quantification of angles (°) between Sz and sSD1 (12 events, 126.9 ± 17.87), Sz and sSD2 (11 events, 143.4 ± 10.77), and sSD1 and sSD2 (12 events, 37.23 ± 8.65), mixed effects analysis with Tukey’s test (F[1.396, 13.96]=26.35; Sz-sSD1 vs. Sz-sSD2 p=0.3938, Sz-sSD1 vs. sSD1-2 p=0.0020, Sz-sSD2 vs. sSD1-2 p<0.0001). For entire fig.: Plotted error bars represent s.e.m., all given ± denote s.e.m.. Depiction of violin plots: median (solid lines), quartiles (dotted lines). Depiction of statistical significance: n.s. not significant, *p < 0.05, **p < 0.01, ***p < 0.001, ****p < 0.0001.

For electrophysiological reference of optically recorded network dynamics, we combined 2p-imaging with local field potential (LFP) recordings (Fig. 1 A, insulated tungsten Ø ∼125 µm). Notably, both in bi-hippocampal wireless LFP recordings in encephalitic mice (Suppl. Fig. 2) and all combined imaging-LFP recordings in CA1 (Fig. 1 E), focal electrographic Sz were faithfully detected bilaterally. Thus, for practical reasons, in experiments involving both chronic in vivo 2p-imaging and LFP recordings, we placed one LFP electrode at the stereotaxic site of hippocampal TMEV inoculation (from Bregma: AP 1.9, ML 1.6, DV 1.5 mm from pial surface), while imaging was performed in contralateral CA1 or CTX (Fig. 1 A, E). In our framework, electrographic and optical hippocampal Sz corresponded well, while neocortical Sz invasion occurred with a delay of seconds (Fig. 1 E).

### Consistent association of seizures with spreading depolarization in hippocampus

Across a total of 205 imaging hours (CA1: 118 hrs; CTX: 87 hrs) in 26 mice and 158 days of viral encephalitis, epileptiform network activity was detected in all mice. In 7 out of those 26 mice, we successfully captured seizures (13 Sz in CA1, 4 mice [Fig. 1 F, blue/left], 8 Sz in CTX, 3 mice [Fig. 1 F, green/right]). Strikingly, imaged hippocampal Sz were consistently followed by a massive second, often multiplexed wave of Ca^2+^ signal, in what appeared to be spreading depolarization / depression (SD, Fig. 1 F, G and suppl. movie 1)^54^. In CA1, neither Sz nor SD ever occurred alone. By contrast, in CTX either isolated SD or Sz-related SD were never detected (Fig. 1 F, G, suppl. movie 2). As described previously, maximum hippocampal Ca^2+^ signal amplitudes of Sz-related SD (sSD) were consistently larger than preceding Sz (Fig. 1 H)^7^. Intriguingly, despite the clear discrepancy of sSD occurrence in CA1 versus CTX, both territories displayed reduced basic Ca^2+^ fluorescence post-sSD (CA1) or post-Sz (CTX) in comparison to the pre-ictal period (Fig. 1 I). However, recovery times to pre-ictal basic Ca^2+^ fluorescence in CA1 were longer than in CTX, on the order of minutes to tens of minutes (Fig. 1 J).

### Hippocampal seizures and sSD propagate in opposite directions

To evaluate potentially conserved micro-progression patterns in naturally occurring hippocampal Sz and sSD during encephalitis, we analyzed spatiotemporal Sz and sSD trajectories across successive events similarly to previous reports^55,56^. Based on the recruitment timepoints of identified individual neurons for a given Sz or sSD, we calculated interpolated linear spatial trajectories of every Sz or sSD across events in all mice with at least three recorded events where Sz and sSD onsets were fully captured (3 mice, CA1, 3.3 ± 0.3 events SEM, Fig. 1 F). As sSD often occurred in two waves immediately following one another, we focused on the trajectory of the first wave (sSD1). Along the antero-posterior and medio-lateral dimension of the imaged field of view (FOV), this resulted in a number of spatial vectors per Sz and sSD1 whose respective standard deviations could be derived and compared to randomized surrogate distributions (for details, see methods). In all three mice, Sz trajectories displayed significantly smaller variability in the observed dataset than would be expected by chance (Fig. 1 K left) suggesting repeated spatiotemporal Sz progression pathways^55–57^. The same held true for 2 out of 3 mice with regards to sSD1 (Fig. 1 K right).

Next, we investigated the relative spatial relationship between individual Sz and successive sSD in CA1 by calculating the angles between spatial trajectories of Sz and sSD1, and sSD1 and sSD2 (Fig. 1 L). Unexpectedly, we found vast angles between Sz and sSD1 trajectories (Fig. 1 L, M). Typically, Sz progressed in a medio-lateral direction (Fig. 1 L left), while sSD1 travelled the opposite way (4 mice, 12 Sz-sSD1 events, median angle 162.01° [range 42.1 – 177.2°]) (Fig. 1 L left/middle, M). By contrast, sSD1 and sSD2 displayed more similar spatial trajectories (Fig. 1 L right, M; 4 mice, 11 sSD1-sSD2 events, median angle 42.5° [0.1 – 100.5°]).

Together, these experiments identified a complete association of naturally occurring Sz and sSD in CA1 during viral encephalitis, whereas this association was not found in CTX. Regardless, both CA1 and CTX displayed reduced basic fluorescence post-sSD or post-Sz, but the post-sSD recovery time of the neuronal network in CA1 clearly outlasted the post-Sz recovery time in CTX. All Sz, and a majority of sSD1 showed significantly non-random spatiotemporal progression, although strikingly, hippocampal Sz and sSD1 travelled in vastly different directions.

### *Post-ictal wandering* is associated with sSD in hippocampus

In the TMEV model, we went on to correlate the observed network dynamics to clinical signs (semiology) related to Sz or sSD. Importantly, during 2p-imaging of focal Sz in awake head-restrained mice, generalized tonic-clonic convulsions never occurred.

What stood out at the clinical level in encephalitis-related seizures was that the mice regularly started locomoting upon optical sSD appearance, which then lasted for minutes (Fig. 2 A, Suppl. Fig. 3). Matching the typical time course of SD (Fig. 1 E, J), in comparison to the pre-ictal period, locomotion was indeed systematically increased post-sSD (pre-ictal 5min vs. post-sSD 5min, Fig. 2 B), fitting well with so-called *post-ictal* wandering, a prominent post-ictal symptom encountered in clinical epileptology. *Post-ictal wandering* is most frequently observed in patients with TLE^1^, and characterized by confusion and unaware wandering around (e.g. on the street). It typically lasts for minutes, at times tens of minutes, with potentially life-threatening consequences. Aside from locomotion time, we also quantified the travelled distance, number of locomotion episodes or maximum locomotion speed, all of which were increased over the 5 min post-sSD period (Fig. 2 C-E). Thus, these experiments showed that during viral encephalitis, hippocampal sSD is associated with the onset of a locomotor phenotype that lasts for minutes, providing a candidate mechanism for the clinical symptom *post-ictal wandering*.

**Fig. 2.**
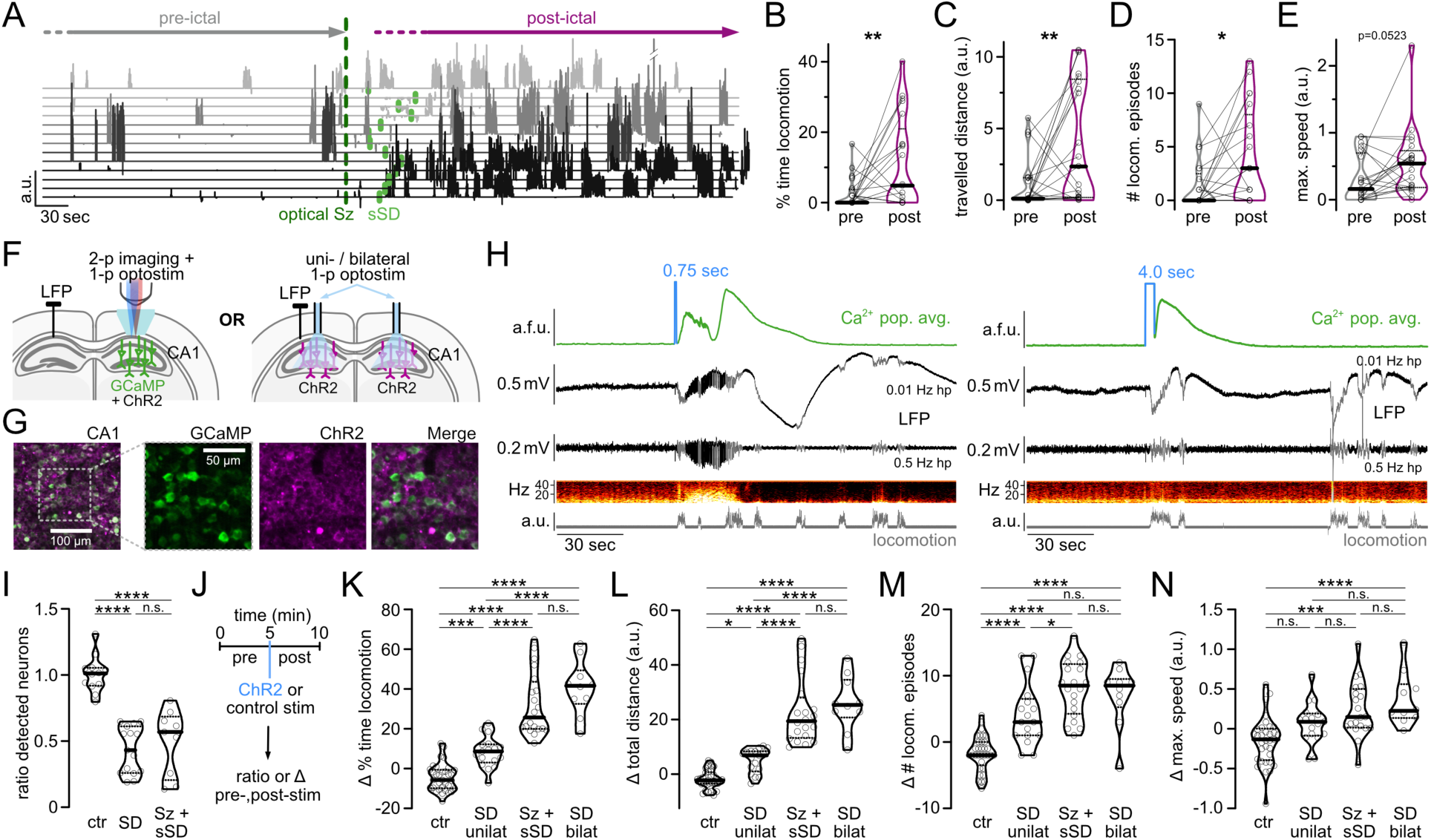
Hippocampal SD triggers increased locomotion. **A**, Locomotion on linear treadmill across 13 Sz-sSD events in all 4 CA1-imaged mice (gray shades) during encephalitis, aligned by optical Sz invasion (dotted green line), sSD onsets marked in light green. **B-E**: Comparison of 21 pre-vs. post-ictal periods (5min each) in all imaged 7 mice. **B**, % time locomotion, paired t-test (df=20; 2.525 ± 0.957 [pre] vs. 11.06 ± 2.83 [post], p=0.0059). **C**, Travelled distance (a.u.: arbitrary units), paired t-test (df=20; 1.055 ± 0.387 [pre] vs. 4.03 ± 0.915 [post], p=0.0041). **D**, # of locomotion episodes, paired t-test (df=20; 1.429 ± 0.519 [pre] vs. 4.0 ± 0.974 [post], p=0.012). **E**, maximum speed (a.u.), paired t-test (df=20; 0.312 ± 0.077 [pre] vs. 0.553 ± 0.11 [post], p=0.0523). **F**, Optogenetic experimental setup. Left (2p/1p/LFP approach): CA1 window for 2p imaging (GCaMP6s or jRGECO1a) and 1p optogenetic stimulation (ChR2), both contralateral to LFP electrode (black pin, at CA1). Right (1p/LFP approach): uni-/bilateral optogenetic stimulation (ChR2) via optical cannulae (CA1) and LFP electrode (black pin, at CA1). **G**, Co-expression of GCaMP6s (transgenic, green) and ChR2 (AAV, magenta) in CA1 (str. pyr.). **H**, Two examples for 2p/1p/LFP approach, Left: 0.75 sec optogenetic stimulation (488 nm) elicits Sz-sSD (optical ipsilateral, electrographic contralateral). Top trace depicts avg pop. Ca^2+^ signal, middle traces LFP (0.01 / 0.5 HP filter) and corresponding spectrogram, bottom trace locomotion; locomotion artifacts in LFP are marked gray in LFP traces; Right: same setting, but 4 sec optogenetic stimulation triggers isolated SD in imaged field of view. In contralat. LFP, neither Sz nor SD are detected. Locomotion artefacts marked in gray in LFP traces. **I**, Ratio of detectable neurons in imaged FOV (CA1) across pre-and post-ictal periods (5min each) in 5 mice, pooled sample #: 12 ctr stim. [561/590nm], 15 SD, 9 Sz-sSD, pre/post ratio: 1.006 ± 0.0396 (ctr), 0.433 ± 0.046 (SD), 49.33 ± 0.083 (Sz-sSD). One-way ANOVA with Tukey’s test (F [2, 33] = 35.37; ctr vs SD p<0.0001, ctr vs Sz-sSD p<0.0001, SD vs. Sz-sSD p=0.7265. **J**, Optogenetic stim. framework for analysis in K to N. **K-N**, Comparison of respective value ∆ between pre-and post-ictal periods (5min each) in 7 mice (2p/1p/LFP or 1p/LFP approach), pooled sample #: 33 ctr stim., 17 unilateral SD, 20 Sz-sSD, 10 bilateral SD. **K**, ∆ % time locomotion: -4,403 ± 1.187 (ctr), 8.331 ± 1.861 (unilat. SD), 32.18 ± 3.540 (Sz-sSD), 40.68 ± 4.325 (bilat. SD). One-way ANOVA with Tukey’s test (F[3,76]=72.44; ctr vs unilat. SD p=0.001, ctr vs Sz-sSD p<0.0001, ctr vs. bilat. SD p<0.0001, unilat. SD vs. Sz-sSD p<0.0001, unilat. SD vs. bilat. SD p<0.0001, Sz-sSD vs. bilat. SD p=0.1862). **L**, ∆ travelled distance (a.u.): -1.805 ± 0.591 (ctr), 5.027 ± 1.063 (unilat. SD), 22.80 ± 2.713 (Sz-sSD), 25.95 ± 3.195 (bilat. SD). One-way ANOVA with Tukey’s test (F[3, 76]=62.57; ctr vs unilat. SD p=0.0178, ctr vs Sz-sSD p<0.0001, ctr vs. bilat. SD p<0.0001, unilat. SD vs. Sz-sSD p<0.0001, unilat. SD vs. bilat. SD p<0.0001, Sz-sSD vs. bilat. SD p=0.7077). **M**, ∆ # of locomotion episodes: -1.697 ± 0.4636 (ctr), 4.176 ± 1.129 (unilat. SD), 8.05 ± 0.95 (Sz-sSD), 7.0 ± 1.461 (bilat. SD). One-way ANOVA with Tukey’s test (F[3, 76]=32.48; ctr vs unilat. SD p<0.0001, ctr vs Sz-sSD p<0.0001, ctr vs. bilat. SD p<0.0001, unilat. SD vs. Sz-sSD p=0.0152, unilat. SD vs. bilat. SD p=0.2561, Sz-sSD vs. bilat. SD p=0.8929). **N**, ∆ maximum speed (a.u.): -0.1516 ± 0.0523 (ctr), 0.072 ± 0.063 (unilat. SD), 0.243 ± 0.077 (Sz-sSD), 0.366 ± 0.106 (bilat. SD). One-way ANOVA with Tukey’s test (F[3, 76]=10.80; ctr vs unilat. SD p=0.0803, ctr vs Sz-sSD p=0.0001, ctr vs. bilat. SD p<0.0001, unilat. SD vs. Sz-sSD p=0.3423, unilat. SD vs. bilat. SD p=0.0876, Sz-sSD vs. bilat. SD p=0.7322). For entire fig.: All given ± denote s.e.m.. Depiction of violin plots: median (solid lines), quartiles (dotted lines). Depiction of statistical significance: n.s. not significant, *p < 0.05, **p < 0.01, ***p < 0.001, ****p < 0.0001.

### Optogenetic dissection of the role of hippocampal SD in *post-ictal wandering*

Although hippocampal sSD appearance coincided with the onset of sustained locomotion during encephalitis, the complete association of hippocampal Sz and sSD precluded us from decisively differentiating the role of Sz versus SD in *post-ictal wandering*. To arrive at a more mechanistic understanding, based on previous studies^58,59^, we established a combined 2p-imaging (GCaMP6s [JAX 025776], or jRGECO [AAV2/1-NES-jRGECO1a] Vector Core Uni Bonn), 1p optogenetics (ChR2, AAV-hSyn-hChR2[H134R]-mCherry, Addgene ID 26976-AAV5) and LFP approach (Fig. 2 F left; 2 G, suppl. Fig 4) that allowed us to generate either Sz or SD in a controlled manner. Using this 2p/1p/LFP approach, Sz or SD were never elicited by 2p-imaging (16X Nikon 0.80 NA, wavelength range 940-980nm [GCaMP] or 1050nm [jRGECO]).

In CA1, Sz could be reliably elicited by a brief square wave light pulse (typically 500-750ms, Coherent Inc., CA, Sapphire LP CW laser, 488nm, power at brain surface 4-5mW/mm^2^) in the imaged FOV (Fig. 2 H). Interestingly, these optogenetically induced Sz in the healthy hippocampus were again consistently followed by sSD, and Sz-sSD could be bilaterally detected (Fig. 2 H left, suppl. movie 3). To reliably trigger isolated SD without a preceding focal Sz, a prolonged light pulse (∼4 sec, 488nm, 4-5mW/mm^2^) was applied, similarly to previous reports (Fig. 2 H right, suppl. Fig. 4 C, suppl. movies 4, 5)^42,43^. Neither Sz nor SD could be elicited by control illumination at 561nm (Coherent Inc., Santa Clara, CA, Sapphire LP CW laser, 7 mW/mm^2^), ensuring that the effects were not unselectively driven by laser light alone. Notably, a comparison of encephalitic hippocampal sSD and optogenetic hippocampal SD with respect to optical signal amplitude, SD_depol_ half-width, and SD progression speed, showed no significant differences (suppl. Fig. 5 A-C). Thus, basic SD features were preserved across our different Sz models. In line with our imaging experiments during encephalitis, both optogenetically triggered Sz-sSD and isolated SD led to an equally strong post-ictal reduction of detectable neurons in the FOV (Fig. 2 I) underscoring functional depression of the CA1 network due to SD. To make sure that these optical imaging results were not confounded by GFP-quenching due to SD-related pH shifts (ongoing neuronal firing in the quenched absence of optical signals), we repeated the optogenetic SD experiments with electrophysiology, now employing hippocampal tetrode recordings (+ optical cannula, see methods) instead of 2p-imaging. In keeping with the imaging experiments, we found a prolonged and profound cessation of firing of a large majority of units upon optogenetic SD in CA1 (suppl. Fig. 5 D).

As unilateral optogenetic Sz induction consistently triggered bi-hippocampal Sz (Fig. 2 H left), we additionally implemented a 1p optogenetic stimulation approach enabling uni-or bilateral hippocampal Sz or SD induction via implanted optical cannulas, combined with LFP recordings (1p/LFP approach; Fig. 2 F right). Here, for optogenetic Sz or SD induction versus control, instead of lasers, fiber-coupled LEDs were employed using the same stimulation durations as mentioned above (Chrolis/Thorlabs: 475 nm or 590nm [2-3mW/mm^2^]). Based on the 2p/1p/LFP or 1p/LFP approach, we then compared a pre-and post-stimulation (for control and SD) or pre-stimulation and post-sSD (for Sz-sSD) period (5min each, Fig. 2 J) analyzing the same locomotor parameters as in the context of encephalitic Sz-sSD (Fig. 2 B-E).

In the initial 2p/1p/LFP experiment, we unilaterally triggered SD or Sz versus unilateral control (Suppl. Fig. 6 A-D). In the 1p/LFP experiment, we carried out bilateral SD, unilateral Sz (except in one animal, where bilateral stimulation was necessary to trigger hippocampal Sz), and uni-or bilateral control stimulations (Suppl. Fig. 6 E-H). Since with respect to all analyzed clinical parameters, uni-versus bilateral control stimulations showed no significant differences, neither within the 1p/LFP configuration (Suppl. Fig. 6 E-H) nor across experimental configurations (Suppl. Fig. 6 I-L), these data were pooled. As the same held true for uni-and bilateral hippocampal Sz inductions within and across experiments, these data were also pooled (Suppl. Fig. 6 I-L, statistics for suppl. Fig. 6 are displayed in suppl. Table 1 and 2).

Remarkably, in comparison to control, isolated optogenetic unilateral SD in CA1 was sufficient to reproduce the locomotor phenotype that we had observed upon Sz-sSD in CA1 during encephalitis, proving that hippocampal SD alone is sufficient to trigger *post-ictal wandering* (Fig. 2 K-N, suppl. Fig. 7). Interestingly, across most tested locomotor parameters (% time of locomotion, travelled distance, number of locomotion episodes), the effect triggered by unilateral hippocampal optogenetic SD stimulation was significantly surpassed by unilateral hippocampal optogenetic Sz-sSD stimulation (Fig. 2 K-N, suppl. Fig. 7). We hypothesized that this difference in effect size could be due to the bilateral hippocampal involvement during focal onset hippocampal seizures, which we had observed during encephalitis and optogenetic experiments (Fig. 1 E, 2 H left). In this case, even a unilaterally triggered hippocampal Sz would, due to its bi-hippocampal recruitment, lead to bilateral hippocampal sSD (Fig. 2 H left). In turn, unlike hippocampal Sz-sSD, due to the different, non-synaptic nearest-neighbor progression of SD, unilaterally triggered SD would typically remain unilateral (Fig. 2 H right). In line with this rationale, uni-or bilateral hippocampal Sz-sSD stimulation produced a similar locomotor phenotype (Suppl. Fig. 6 E-H), and Sz-sSD stimulation systematically produced a stronger phenotype as compared to unilateral optogenetic SD stimulation, except for maximum speed (Fig. 2 K-N, suppl. Fig. 6 A-D and I-L). Yet crucially, Sz-sSD and bilateral SD stimulation consistently evoked a similar clinical phenotype across analyzed locomotor parameters (Fig. 2 K-N, suppl. Fig. 6 E-L, suppl. Fig. 7). Together, these experiments unveiled i) a mechanistic role of hippocampal SD in *post-ictal wandering*, ii) a scaling of severity of the clinical phenotype depending on uni-versus bilateral hippocampal SD, and iii) similar clinical effect sizes across hippocampal Sz-sSD and bilateral SD. Therefore, rather than Sz themselves, these experiments point towards a role of hippocampal SD in *post-ictal wandering*.

### Optogenetic induction of hippocampal SD and neocortical widefield imaging

While we had never observed Sz-sSD in motor cortex using 2p-imaging during encephalitis, our imaged FOV was confined to ∼700x700 µm. Thus next, with the optogenetic approach of on-demand focal SD induction at hand, and combined LFP recordings and one-photon (1p) widefield Ca^2+^ imaging, we went on to investigate the potential spread of optogenetic unilateral hippocampal SD to CTX^28,29^. For hemispheric neocortical 1p-imaging (30 Hz scanning, 470 / 405 nm, LED M470L3 / M405L3, Thorlabs) through cleared skull in awake adult mice, we employed an inverted tandem lens macroscope^60^, and transgenic tetO-GCaMP6s / CaMK2a-tTA mice (JAX 024742 / 007004) expressing GCaMP6s in excitatory neurons across all cortical layers (for details, see methods) (Fig. 3 A).

**Fig. 3.**
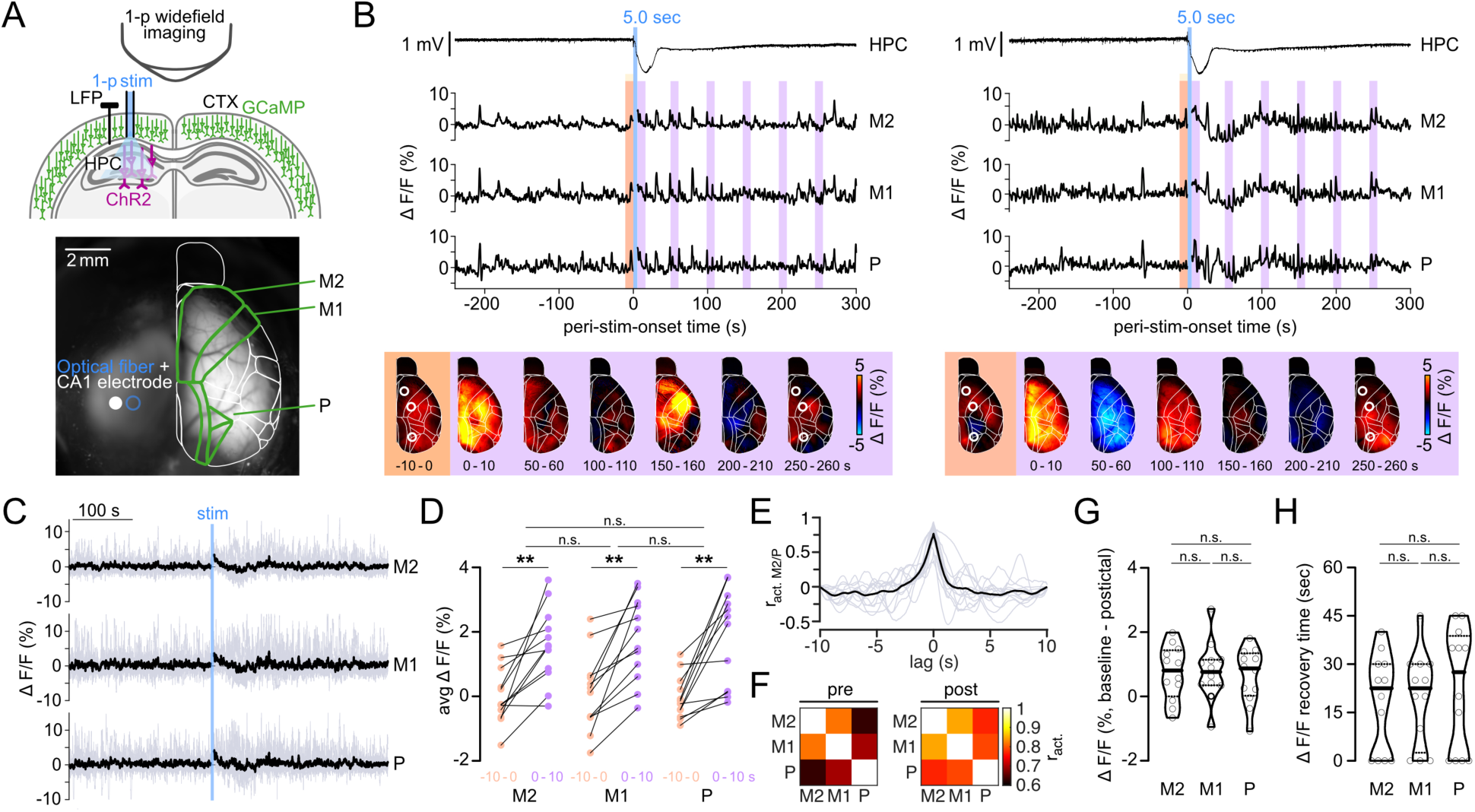
Hippocampal SD coactivates but does not invade neocortex. **A**, Top: Experimental setup, widefield Ca^2+^ imaging of neocortex (CTX) in tetO-GCaMP6s/CaMK2a-tTA mice through cleared skull, in combination with hippocampal optogenetic 1p stimulation (ChR2, optical cannula at CA1) and LFP (black pin, at CA1), Bottom: Example image of cortical surface after skull clearing, and implant sites of LFP electrode (white filled circle) and optical fiber (blue circle). Overlaid white lines show Allen CCF borders. **B**, Two examples of stimulus-related CTX activity. Hippocampal (HPC) LFP of optogenetic SD stimulation (5 sec, blue line), and cortical widefield Ca^2+^ activity in secondary/primary motor (M2, M1) and parietal cortex (P). Corresponding to pre-(orange) and post-stim. periods (violet) 10-sec avg ∆ F/F images of CTX activity (bottom, white circles depict M2, M1, P). Note immediate post-stim. widefield CTX activation and continued post-stim. activity, either without any baseline F shift (left) or a brief transient negative baseline shift (right), yet always with continued presence of cortical Ca^2+^ transients. **C**, Superposition of peri-stim. CTX ∆ F/F activity traces (individual traces in gray, mean in black; 3 mice, 12 HPC SD stim. [4/mouse, 1/day]). **D**, Quantification of pre-(-10 -0 sec to stim., orange) vs. post-stim. (0 – 10 sec from stim., violet) avg CTX activity in M2 (0.01 ± 0.252 [pre] vs. 1.589 ± 0.337 [post]), M1 (0.118 ± 0.348 [pre] vs. 1.778 ± 0.371 [post]) and P (-0.044 ± 0.196 [pre] vs. 1.831 ± 0.432 [post]). One-way ANOVA with Šidák’s test (F[2.146, 23.60] = 17.98; pre vs. post: M2 p=0.0029, M1 p=0.0014, P p=0.0018). No significant differences of ∆ (post – pre) activity among M2, M1, and P. One-way ANOVA with Tukey’s test (F[1.837, 20.21]=0.5231; M2 vs. M1 p=0.946, M2 vs. P p=0.657, M1 vs. P p=0.766). **E**, Cross-correlation of peri-stim. activity of M2 and P (12 HPC SD stim., [4/mouse, 1/day], individual traces in gray, mean in black), mean lag = 0 sec. **F**, Pre-and post-ictal (5min each) activity correlation of M2, M1, and P. Note increased correlation of activity over 5min post HPC SD stimulation. **G**, CTX pre-vs. post-ictal minimal F (∆ min F baseline/min F post-ictal; M2 0.739 ± 0.243, M1 0.781 ± 0.26, P 0.643 ± 0.247), One-way ANOVA with Tukey’s test (F[2, 33]=0.0807; M2 vs. M1 p=0.992, M2 vs. P p=0.96, M1 vs. P p=0.919). Note that all areas show a decreased post-stim. min F on avg. **H**, Comparison of cross-area F recovery time (sec, M2 18.33 ± 4.323, M1 19.58 ± 4.15, P 22.5 ± 5.418), One-way ANOVA with Tukey’s test (F[2, 33]=0.2102; M2 vs. M1 p=0.98, M2 vs. P p=0.804, M1 vs. P p=0.898). For entire fig.: All given ± denote s.e.m.. Depiction of violin plots: median (solid lines), quartiles (dotted lines). Depiction of statistical significance: n.s. not significant, *p < 0.05, **p < 0.01, ***p < 0.001, ****p < 0.0001.

Around six weeks before the actual experiment, pAAV-hSyn-hChR2(H134R)-mCherry was stereotactically injected into the left hippocampal CA1 region (coordinates from Bregma: AP -1.9, ML -1.6, DV -1.5 mm). In addition, a custom-made optrode comprising an insulated tungsten electrode (Ø ∼125 µm) and an optical fiber (Ø 200 µm, NA = 0.37) were chronically implanted above the left hippocampus, for LFP recordings and optical stimulation.

Similar to our previous optogenetic experiments, hippocampal SD could be reliably induced by a square wave light pulse (5 sec, 488 nm, 15 mW/mm^2^, 1p CW laser READYBeamÔ Bio2, FISBA AG, SUI) (Fig. 3 B). For practical procedural reasons related to the optrode implant, imaging was carried out in the cortical hemisphere contralateral to hippocampal SD induction (Fig. 3 A). While a widespread coactivation of CTX (e.g. motor cortex: M2, M1; parietal cortex: P) was observed during optogenetically induced hippocampal SD across 12 recordings (4 mice, 3 SD each, one SD stim/d, Fig. 3 B-D), no SD or wave-like invasion resembling SD dynamics were ever observed in CTX. Moreover, no activation lags could be found across distant neocortical areas (e.g. M2/P, Fig. 3 E), and CTX continued to show typical transients of Ca^2+^ activity (Fig. 3 B). Notably, neocortical areas showed an enhanced correlation of activity in the post-stimulation period (Fig. 3 F). Still, similar to our 2p-imaging results in CTX during encephalitis, cortical dynamics across regions also often displayed, although moderately, a reduction of basic fluorescence in the immediate post-stimulation period despite the absence of SD (Fig. 3 B right, 3 C and G, suppl. Fig. 8). Importantly, also during this time, Ca^2+^ transients continued without interruption (Fig. 3 B right). Further, as in our 2p-imaging experiments, the recovery to pre-stimulation basic fluorescence was consistently fast in CTX (<1 min, Fig. 3 H). Together, these experiments recapitulated results from our 2p-imaging experiments, and corroborated the evidence that hippocampal SD typically does not propagate to CTX. Moreover, neocortical Ca^2+^ transients appeared to continue throughout the post-stimulation period without signs for activity depression, and, in contrast to SD dynamics, showed a rapid recovery from reduced basic fluorescence to pre-stimulation conditions.

### Seizure-related focal SD in human epilepsy

The current EEG-standard with a bandwidth of 0.5-70 Hz^61^ renders SD invisible in clinical practice (suppl. Fig. 09). While SD related to spontaneous Sz has been documented in rodent epilepsy models^29^, it is not known whether it occurs in human epilepsy. Thus, we finally sought to examine the existence, and potentially regional preference of sSD in human focal epilepsy through multi-regional Behnke-Fried (BF) electrode recordings, in the context of stereotactic depth macro-electrode recordings (SEEG) during pre-epilepsy-surgery diagnostic work-up (Fig. 4 A, B, see methods for details).

**Fig. 4.**
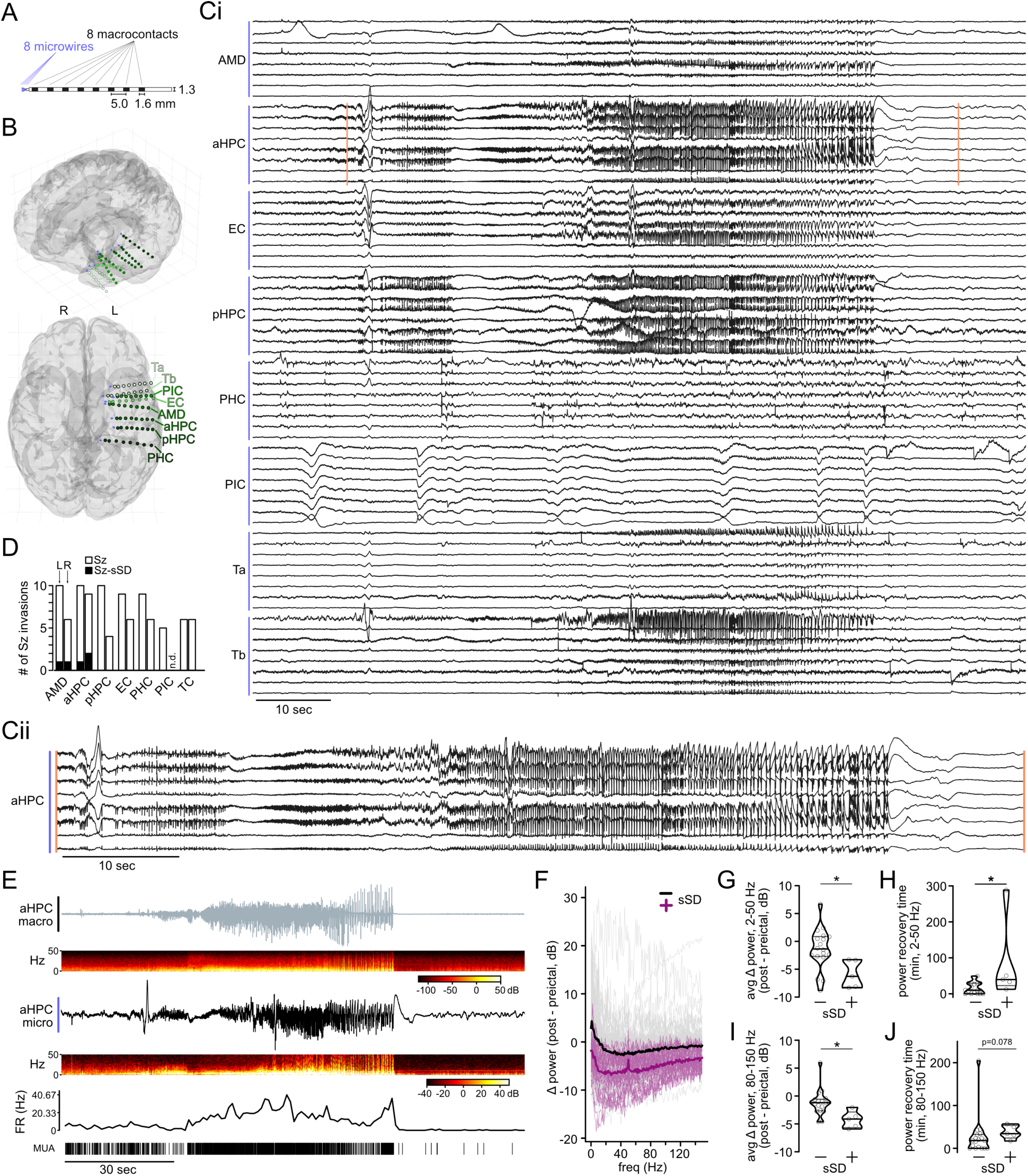
Focal Sz-sSD occurs in human epilepsy. **A**, Scheme and dimensions of individual Behnke-Fried (BF) electrodes composed of depth-electrode macro-contacts and microwire bundle. **B**, 3D-reconstructed brain MRI and location of implanted BF electrodes. Green dots indicate macro-contact locations, shades of green designate different brain regions and violet markers depict microwire bundles. AMD: amygdala, aHPC: anterior hippocampus, EC: entorhinal cortex, pHPC: posterior hippocampus, PHC: parahippocampal cortex, PIC: piriform cortex, Ta: temporal cortex a, Tb: temporal cortex b. **C i**, LFP traces from BF microwire bundles across recorded regions (0.1 HP filter, referenced to local common average) of a temporomesial focal onset Sz. **C ii**, magnified inset from Ci, aHPC (as marked by orange lines). Note slow LFP shift at the end of Sz, confined to HPC. **D**, Regional occurrence count of sSD across all Sz-invaded regions (all Sz/patients included). **E**, aHPC, most distal macro contact trace (top), corresponding microwire trace (middle) and multi-unit activity from the same microwire (bottom). **F**, Change in power between the first postictal minute and the preictal minute. Each line represents one Sz-invaded microwire, color shades indicate Sz-invaded (gray, sSD-neg.) and Sz-sSD-invaded (purple, sSD-pos.) regions, bold lines depict the mean of each group. **G**, Comparison of respective ∆ pre-and post-ictal broadband power (dB, 2-50 Hz) in sSD-neg. (18 events) vs. sSD-pos. (5 events) Sz-invaded regions, unpaired t-test (-1.365 ± 0.8728 [sSD-neg.] vs. -5.870 ± 1.107 [sSD-pos.], p=0.0189). **H**, Comparison of postictal broadband power recovery time (min) of sSD-neg. (18 events) vs. sSD-pos. (5 events) Sz-invaded regions, Mann-Whitney test (16.7 ± 3.84 [sSD-neg.] vs. 84.87 ± 51.29 [sSD-pos.], p=0.0112). **I**, Comparison of respective ∆ pre-and post-ictal high-gamma power (dB, 80-150 Hz) in sSD-neg. (18 events) vs. sSD-pos. (5 events) Sz-invaded regions, unpaired t-test (-1.184 ± 0.544 [sSD-neg.] vs. -4.227 ± 0.6825 [sSD-pos.], p=0.0117). **J**, Comparison of postictal high-gamma power recovery time (min) of sSD-neg. (18 events) vs. sSD-pos. (5 events) Sz-invaded regions, Mann-Whitney test (27.05 ± 11.00 [sSD-neg.] vs. 37.97 ± 7.08 [sSD-pos.], p=0.078). For entire fig.: All given ± denote s.e.m.. Depiction of violin plots: median (solid lines), quartiles (dotted lines). Depiction of statistical significance: n.s. not significant, *p < 0.05, **p < 0.01.

As an initial cohort, we included four patients (2 female / 2 male patients, age range 23-50 yrs) with refractory focal epilepsy based on different pathologies, a history of post-ictal symptoms comprising post-ictal wandering, aphasia or confusion, and depth electrodes in the temporal lobe (Fig. 4 B, for implant schemes and clinical information, see suppl. Fig. 10). Across a total of 272 BF microelectrodes (8/bundle), 7 brain regions (amygdala, hippocampus [ant., post.], entorhinal / parahippocampal / piriform / temporal cortex), and 39 recording days during pre-surgical evaluation, 16 focal-onset seizures were recorded. Strikingly, we found what appeared to be highly localized sSD in the BF microwire recordings (Fig. 4 Ci, Cii) in every patient (4/16 Sz in total). In the per-region analysis of Sz-invasions, in line with our murine recordings, we found sSD to occur primarily in temporomesial regions, while it was not detected in the other recorded regions (Fig. 4 C, D). At the same time, sSD was not visible at the clinical macroelectrodes (Fig. 4 E, top panel). On average, both non-sSD and sSD regions displayed a postictal/post-sSD reduction in spectral power (Fig. 4 F), but sSD regions showed a more profound decrease, and a delayed recovery to the pre-ictal condition (Fig. 4 G, H). Consistent with our murine results and previous literature^8,17,32,62^, at the neuronal unit level, sSD invasion coincided with a break-down of neuronal firing (Fig. 4 E, middle and bottom panel). To compare neuronal activity independently of stably trackable neuronal units, we used high gamma (80-150 Hz) oscillatory power as a proxy, as previously described^63^. Again, on average, both sSD-neg. and sSD-pos. regions showed a decreased post-ictal high gamma power, but sSD-pos. regions displayed a stronger reduction, and delayed recovery (Fig. 4 I, J). In sum, these recordings unveiled that focal sSD exists in human focal epilepsy, and suggest an increased propensity of temporomesial regions to experience sSD across a cohort of 4 patients. Further, among Sz-invaded areas, sSD-pos. regions take longer to recover to their pre-ictal baseline in comparison to sSD-neg. regions.

## Discussion

This work suggests that seizure-associated focal spreading depolarization (sSD) is a pathoclinical hallmark of epilepsy. We find profound region-specific differences in sSD occurrence, characterize its progression, and show that temporomesial SD causes post-ictal wandering. Then, in a cohort of 4 human epilepsy patients with BF electrode recordings, we show that sSD exists in human focal epilepsy. In this cohort, sSD displayed a similar temporomesial propensity as in our murine experimental models.

While we observed sSD primarily in temporomesial structures in both mice and humans, several aspects require discussion in this regard. First, our analyses were based on focal onset seizures with known or suspected origin in the temporal lobe, with known or suspected seizure onset zones (SOZ) situated there too. Therefore, sSD emergence may simply depend on its spatial relationship to SOZ and (peri-)lesional tissue. Still, there are basic neuroanatomical features that inherently predispose e.g. the hippocampus to sSD. First, hippocampal wiring with itself and other regions renders it more excitable than other portions of the brain such as the neocortex^64,65^. In line with this longstanding notion, our hippocampal optogenetic Sz-induction protocol failed to induce seizures in healthy neocortex. At the synaptic level, the CA3 region contains so-called “conditional detonator” synapses that can strongly drive postsynaptic firing^66^, which may be why CA3 displayed a high propensity for SD upon repetitive mossy fiber stimulation under hyperexcitable conditions^67^. Further, aside from its neuronal connectivity, recent data suggests that the hippocampus displays relatively reduced blood oxygenation and neurovascular coupling in comparison to neocortex due to differences in vascular architecture^68^. Among other potential contributing factors such as subregion-specific oxidative metabolic capacity^69^, this may make the hippocampus susceptible to an energy shortage during seizures, promoting sSD. We propose that in addition to possible (peri-)lesional vulnerabilities for sSD, basic neuroanatomical differences across brain regions support differential sSD occurrence rates. This carries implications for notions revolving around regional post-ictal EEG signal depression and its spatial overlap with the assumed SOZ^70^. It is possible that e.g. the hippocampus, even if only secondarily recruited into a Sz, may be at risk for sSD due to its anatomy. This could then prompt post-ictal signal depression in this territory without it being part of the SOZ.

Adhering to the neuroanatomical argument, regarding the disparate spatiotemporal trajectories of Sz and sSD in our imaging experiments, work by Scharfman on SD susceptibilities of hippocampal sub-regions^67^ may provide a potential explanation for this unexpected result. We always imaged CA1 contralateral to the TMEV CA1 injection site, and consistently observed a mediolateral Sz spread. sSD mostly travelled the other way, which points towards CA2 or CA3 as the source of sSD^67^. Based on Scharfman’s dual region electrical recordings, her observation that mossy-fiber-stimulation-triggered SD in CA3 travelled back to dentate gyrus (DG)^67^, and our CA1 imaging results, one could hypothesize that CA3-prompted sSD propagate both towards DG and CA1. Alternatively, CA2 could be an sSD source, as it receives strong excitatory input from entorhinal cortex, and has been shown to display enhanced excitability in chronic epilepsy^71,72^. Finally, it is also conceivable that multiple potential sSD generators exist across CA regions.

A main finding of this study is that hippocampal SD triggers post-ictal wandering. Importantly, this does not *per se* mean that the hippocampus is the primary driver of the observed phenotype. While a transient shutdown of hippocampal networks will contribute to unaware post-ictal wandering e.g. through navigational confusion, it is likely that the enhanced locomotion upon hippocampal SD is brought about by a number of interconnected brain regions^73,74^ that interact with the hippocampus and that may be indirectly or directly affected by hippocampal SD, including the septum^75,76^ and mesencephalic structures^73,77^. Clearly also, while in our murine models, sSD was observed only in hippocampus during viral encephalitis, and optogenetic hippocampal SD did not invade neocortex, this does not mean that it never does (it can, see ^28,29^), nor that it does not invade other subcortical structures (we hypothesize that it does), nor that it does not invade the hippocampus if emerging elsewhere (e.g. in the neocortex^28^).

At a wider scope, this works carries implications for epilepsy research and neurology. Surely, while much has been learned about SD in neurological research e.g. on migraine or traumatic brain injury^10–12^, the relationship between SD and clinical symptoms in epilepsy has remained understudied. Part of the reason is that most studies on SD in epilepsy have been carried out in vitro, or under anesthesia^8,9,23,26,49,50^, which precludes clinical correlation. At other times, it was not primarily intended to link neurophysiological measures to clinical semiology, i.e. in studies focusing on SD-related Sz termination^50,51^. Of note, different types of epilepsies (e.g. acquired vs. genetic epilepsy) and seizures (focal vs. bilateral tonic clonic) will likely affect the occurrence and spatial extent of SD (e.g. local vs. [bi-]hemispheric), and thus impact clinical phenotypes^78^. Remarkably, most epilepsy research studies do not specifically distinguish Sz with or without SD, although SD has long been suggested to represent a ‘separate entity’ (see Bureš et al.^24^, p.10). Based on this suggestion and the results shown here, it may indeed be the case that some of the previously observed effects attributed directly to Sz are instead mediated by sSD. Notably, as Sz and sSD can co-occur at the same time in different brain regions, this also extends to clinical semiology. Some previously described “ictal” symptoms may present compound semiology based on spatially separate, simultaneous Sz and sSD. Re-visiting reported effects and effect-sizes, and clinical symptoms through the lens of Sz versus sSD will likely propel new diagnostic and therapeutic research on many acute and chronic clinical circumstances that involve seizures, and thus potentially, sSD.

Due to the current EEG standard (≥0.5 Hz)^61^, SD is invisible in routine clinical care. As a result, medical professionals regularly encounter post-ictal symptoms, but are blind to SD as a potential major determinant of these phenomena. As there is a traditionally strong focus on the “ictus” in epileptology (highlighted by current terminology), such symptoms are then usually related to Sz, and sSD plays little to no role in clinical care. Inversely, it is known in basic research that sSD terminates ongoing Sz^27,50^, but since SD-mediated Sz termination have been studied mostly in vitro or under anesthesia, the clinical impact of sSD could not be studied. Thus, our results underscore the necessity of a close cross-talk between clinical and basic research disciplines. Clearly, sSD can have potentially life-threatening clinical consequences^4,5^. In our view, caution is required if sSD is proposed as a protective factor of the brain against seizures, and as a potential epilepsy treatment tool^50,52^. Rather, our work supports the notion that SD constitutes a transient homeostatic break-down of a biological system driven beyond its physiological range of operation^32^.

There are of course limitations to this study. First, although our murine findings held true across recording modalities, two different murine models of hippocampal Sz and SD, and under healthy (optogenetics) and disease (TMEV) conditions, we do not know whether these findings extend across many different etiologies. Aside from the factors discussed above, the exact parameters that favor sSD emergence in hippocampal Sz are not yet clear, and may include e.g. Sz duration, extent of spatial Sz invasion and hippocampal subfields, Sz dynamics (e.g. tonic vs. clonic), internal state or transition between states (e.g. wakefulness, sleep). Further, there are limitations related to the human data included in this study. In our wide-bandwidth AC recordings, the lowest possible high-pass filter was 0.1 Hz for technical reasons. Therefore, it is possible that some sSDs were missed, and if anything, the true occurrence rate may be higher. Another reason for a possible underestimation of the true sSD occurrence rate is that current depth electrode recordings strongly spatially undersample targeted brain areas. In the near future this limitation may become softened to some extent, on the heels of recent advances in high-density electrode arrays^79–82^. First steps towards a systematic investigation of sSD in larger patient cohorts will require coordinated efforts by institutions that have access to wide-bandwidth AC or DC-coupled intracranial electrical recordings in epilepsy patients. Beyond the currently available recording modalities in clinical practice, one should consider newly available technology for potential high-fidelity full-bandwidth recordings of Sz, SD and sSD in humans, e.g. biocompatible flexible microtransistors^28^.

In sum, this work sets stage for a wider discussion of the pathoclinical role of sSD in epilepsy, and a potential re-consideration of the clinical EEG filter standard^61^. Beyond post-ictal wandering, our results suggest that focal sSD could underlie other post-ictal symptoms, e.g. confusion, receptive aphasia, navigational impairment, or defensive-like aggression^3,83,84^. Finally, it seems plausible that sSD does not only trigger immediate post-ictal symptoms, but also exerts chronic effects, e.g. on cognitive function. Thus, it may play a role in comorbidities of epilepsy, and other diseases encompassing temporal lobe pathology and seizures, e.g. neurodegenerative diseases, which are not primarily treated by epileptologists.

## Author approvals

All authors have seen and approved the manuscript, and the manuscript has not been accepted or published elsewhere.

## Acknowledgements

We sincerely thank Lea Adenauer for outstanding technical support, and Negar Nikbakth, Nicola Masala, and Martin Pofahl for their technical advice. We appreciate Lisa Rottenfußer’s support and input during her M.Sc. lab rotation. Generally, we are grateful to members of the Wenzel, Beck, Ewell, and Mormann laboratories for comments. This work was supported by the BONFOR Program at University of Bonn (M.W.: #2019-2-04), the Hertie Network of Excellence in Clinical Neuroscience (M.W.: #P1200008), and the German Research Foundation (DFG; SFB1089 to M.W., L.E., H.B., and #504342801 to M.W.). The work was further supported by the iBehave network (M.W., S.M., H.B.), the Helmholtz association (S.M.: VH-NG-1611), and the European Research Council (M.W.: StG #101039945). We also acknowledge the support of the Viral Vector Service Core Facility, and the Imaging Core Facility of the Bonn Technology Campus Life Sciences funded by the Deutsche Forschungsgemeinschaft (DFG, German Research Foundation, project #388169927).

## Author contributions

M.W. conceived the project. M.W. and B.M. wrote the paper with input from all authors. Experimental contributions: Multimodal 2-photon imaging experiments: B.M., M.W., M.K., M.B., T.T., T.O.. Multimodal 1-photon imaging experiments: N.B., S.M.. Tetrode recordings: A.N.H., M.K.. Wireless LFP recordings: A.B., J.P., M.W.. Optogenetics (combined with imaging/electrophysiology, via cranial window/cannulas): M.K., M.W., B.M., T.T., A.N.H., N.B., S.M.. Immunohistochemistry and confocal microscopy: L.K., M.B., Š.G., M.S-S., A.B., J.P.. TMEV was produced and provided by I.G. and W.B.. All data processing and analyses were carried out by B.M. and M.W., except: Multimodal 1-photon imaging experiments: S.M., N.B., M.W.. Tetrode recordings: A.N.H., L.E., A.G.G., M.K.. Immunohistochemistry and confocal microscopy: L.K., M.B., Š.G., M.S-S., A.B., J.P.. Processing and analysis of human recordings: A.G,G., M.W., F.M.. H.B., R.S., and F.M. provided infrastructure, experimental, analytical, and clinical expertise.

