## Supplementary material for "Region-specific spreading depolarization drives aberrant post-ictal behavior": Materials and Methods

### Mitlasóczki et al., Materials and Methods

### Experimental models and subject details

All experiments were performed with care and in compliance with the European, German national, and institutional regulations (animal protocols 81-02.04.2019.A288, 81-02.04.2022.A354). All animals were housed at 12h reverse light cycles (12 hours of darkness from 7am – 7pm, 12 hours of light). Food and water access were provided *ad libitum*. For all experiments described in detail below, except for widefield imaging experiments, adult male and female wild-type C57BL/6J or transgenic Thy1-GCaMP6s mice (C57BL/6J-Tg(Thy1-GCaMP6s)GP4.12Dkim/J; Jackson Laboratories; RRID:IMSR_JAX:025776)[1,2] at postnatal age of 3-6 months were used, and randomly assigned to experimental groups. For neocortical widefield imaging, 4 adult transgenic tetO-GCaMP6s/CaMK2a-tTA male mice at postnatal age of 3-6 months were used, expressing GCaMP6s specifically in excitatory neurons across all cortical layers. Mice were obtained by crossing the two mouse lines DBA-Tg(tetO-GCaMP6s)2Niell/J (RRID:IMSR_JAX:024742), and Cg-Tg(CaMK2α-tTA)1Mmay/DboJ (RRID:IMSR_JAX:007004)[3]. Human depth electrode implantations were carried out solely for clinical reasons, in the context of pre-epilepsy-surgery diagnostic work-up of patients with medically refractory focal epilepsy, at the Dept. of Epileptology at University Medical Center Bonn (for clinical information, see Suppl. Fig. 10). With respect to additional depth BF-microwire recordings, all participants gave their written informed consent, and the study was approved by the Medical Institutional Review Board of the University of Bonn.

### Method details

#### Experimental and surgical procedures

**Viral injections:** Theiler's encephalomyelitis virus (TMEV) model. For multimodal 2p-imaging experiments during encephalitis, we used the recent Theiler's murine encephalomyelitis virus (TMEV) etiocopy mouse model of TLE originally established by Libbey et al.[4]. Local intracerebral injection of TMEV (Picorna-virus, Daniel [DA] strain, obtained from Ingo Gerhauser and Wolfgang Baumgärtner, University of Hannover, Germany) induces self-limiting viral encephalitis (~14 days post injection [p.i.]), results in acute disease stage seizures (Sz, mostly ~2-5 days p.i.) (Fig. 1 C) and pronounced immune activation (Suppl. Fig. 1 A-D)[5,6]. Towards chronic stages, the TMEV model commonly leads to hippocampal sclerosis (Suppl. Fig. 1 E, F) and TLE after a post-infectious period of epileptogenesis[7,8]. For stereotactic injection of TMEV, mice were first anesthetized with isoflurane (initial dose 3-4% partial pressure in air [p.p.a.], then reduction to 1-1.4%). Peri-procedurally (-0.5 to 72 hours), mice received ketoprofen for pain relief (Gabrilen, Mibe; 5 mg/kg body weight [b.w.], subcutaneous [s.c.]). To induce viral encephalitis, 10µl viral solution of TMEV DA strain (~1.27x$10^8$ plaque forming units / ml, in DMEM culture medium; NanoFil 34G beveled needle, used with NanoFil 10µl syringe, both World Precision Instruments) was injected unilaterally into the hippocampal CA1 region (coordinates from bregma: AP = -1.9 mm, ML = ±1.6 mm, DV = 1.5 mm) using a programmable microinjector (World Precision Instruments), through a burr hole established with a dental drill. For injection of control mice, DMEM culture medium only was used. The injection speed was kept at a constant 100 nl/min to avoid tissue displacement due to the injected volume. 10 minutes after completion of injection, the injection needle was slowly withdrawn, and the burr hole closed with dental cement (Tetric EvoFlow).

Over 7 days p.i. (TMEV, or DMEM only), mice underwent continuous video monitoring (with intermittent breaks for multimodal imaging experiments) in order to obtain a basic overview of Sz occurrence throughout viral encephalitis (Fig. 1 C). Recorded videos were manually analyzed for Sz with a Racine stage of 3 and above. Sz observed outside the monitoring setup (e.g. during transfer to the 2p microscope) were documented as well. Sz were only detected in TMEV mice, not in control mice.

Upon completion of experiments (max. 3 months p.i.), mice were deeply anesthetized and perfused with PFA, and their brains collected for subsequent immunohistochemistry (for details, see below in separate section).

Adeno-associated virus (AAV) mediated expression of ChR2 or jRGECO in CA1. AAV-mediated neuronal expression of channelrhodopsin 2 (ChR2) or jRGECO1a in the hippocampal CA1 region was achieved via stereotactic injection (coordinates from bregma: AP = -1.9 mm, ML = ±1.6 mm, DV = 1.5 mm). Thirty minutes after administration of ketoprofen (Gabrilen, Mibe; 5 mg/kg b.w., s.c.) mice were anesthetized with isoflurane (initial dose 3-4% p.p.a., then reduction to 1-1.4%). Eye-ointment was applied (Bepanthen, Bayer), and body temperature was maintained at 37°C by closed-loop regulation on a warming pad (TCAT-2LV, Physitemp). Then, two burr holes, about 1 mm apart and centered around the mentioned coordinates, were established using a dental drill. Using the same type of syringe and needle as for stereotactic TMEV injection, 400 nl of pAAV-Syn-hChR(H134R)-mCherry (AAV5, gift from Karl Deisseroth, Addgene_26976, titer ≥ $7\times10^{12}$ vg/ml,) diluted 1:1 with PBS were injected via each burr hole, in wildtype or transgenic mice listed under section *subject details*. For simultaneous jrGECO1a and ChR2 expression, using the same framework as just described, CAMKIIa-jNES-jRGECO1a (AAV2/1, University of Bonn Vector Core, injection volume 400 nl) and pAAV-Syn-hChR(H134R)-GFP (AAV8, Addgene_58880, titer ≥ $7\times10^{12}$ vg/ml, injection volume 400 nl)[9] were injected in bl6 wildtype mice.

**Head fixation ring implantation.** A head fixation ring (titanium, 2g) was implanted as previously described[10]. Thirty minutes after admission of ketoprofen (Gabrilen, Mibe; 5 mg/kg b.w., s.c.) and buprenorphine (Buprenovet, Bayer; 0.05 mg/kg b.w., 0.1 ml/20g b.w., s.c.) for analgesic treatment, mice were anesthetized with isoflurane (initially 3-4% p.p.a., then reduction to 1-1.4%). Eye-ointment was applied (Bepanthen, Bayer), and body temperature was maintained at 37°C by closed-loop regulation on a warming pad (TCAT-2LV, Physitemp). The hair above the implantation site was removed, the skin disinfected using iodine solution, and 10 % lidocaine applied as a local anesthetic. To expose the skull, a circular piece of scalp (Ø ~6 mm) was cut out, and all connective tissue thoroughly removed using 3 % H2O2 and a scalpel. Then, the exposed skull was coated with two-component dental adhesive (Kerr OptiBond FI) that was UV-cured for 30-60 seconds. Thereafter, the titanium ring was attached using dental cement (Tetric EvoFlow). In mice used for optogenetic experiments, a unilateral injection of ChR2 followed immediately, as described above. Over three days post surgery, mice received 5 mg/kg b.w. ketoprofen s.c. and buprenorphine (Buprenovet, Bayer; 0.05 mg/kg b.w., 0.1 ml/20g b.w., s.c.) as analgesic and anti-inflammatory treatment.

**Cranial window implantation for chronic two-photon imaging.** For multimodal two-photon imaging experiments, a chronic cranial window (hippocampal CA1 or neocortex) was established. 30 minutes prior to induction of anesthesia, buprenorphine was applied for analgesia (Buprenovet, Bayer; 0.05 mg/kg b.w., 0.1 ml/20g b.w., s.c.). Moreover, dexamethasone (Dexa, Jenapharm; 0.1 mg/20 g b.w., i.p.) and ketoprofen (Gabrilen, Mibe; 5 mg/kg b.w., s.c.) were administered to treat inflammation, swelling, and pain. Mice were anesthetized with isoflurane (initially 3-4% p.p.a., then reduction to 1-1.4%). Eye-ointment was applied (Bepanthen, Bayer), and body temperature was maintained at 37°C by closed-loop regulation on a warming pad (TCAT-2LV, Physitemp).

Chronic hippocampal windows were established as previously described[2,10–12]. Briefly, using a dental micromotor drill, a circular craniotomy (Ø ~3 mm) was established above the dorsal hippocampus within the central foramen (Ø ~7 mm) of the implanted titanium ring. Neocortical tissue was carefully aspirated until the alveolar fibers above the CA1 region became visible. Hereupon, a custom-made silicon cone (top Ø 3 mm, bottom Ø 2 mm, depth 1.5 mm, RTV 615, Momentive) attached to a thin cover glass (Ø 5 mm, thickness 0.17 mm) was slowly lowered onto CA1 using a micro-manipulator (Luigs & Neumann), and fixed on the skull around the edges of the cover glass using dental cement (Tetric EvoFlow).

Chronic neocortical windows were established as previously described[13–15]. Similar to the hippocampal window procedure, a circular section of the skull was thinned using a dental micromotor drill, until a small piece of skull (Ø ~3 mm) could be removed effortlessly with fine forceps. The dura mater was kept intact. Finally, a thin layer of low-temperature melting agarose (Sigma-Aldrich) was applied, the craniotomy covered with a thin glass cover (Ø 3 mm, thickness 0.17 mm), and fixed using dental cement (Tetric EvoFlow). Over three days post surgery, mice received 5 mg/kg b.w. ketoprofen s.c. and buprenorphine (Buprenovet, Bayer; 0.05 mg/kg b.w., 0.1 ml/20g b.w., s.c.) as analgesic and anti-inflammatory treatment. Further, during these three days, mice were carefully monitored twice daily. Animals typically recovered from surgery within 24–48 hours, showing normal activity and no signs of pain or distress. At least two weeks of recovery time was allowed before animals were habituated to the respective experimenters and experimental setups, and at least 4 weeks passed postoperatively before multimodal experiments were started.

**Electrode implantation for field electrophysiology in multimodal two-photon imaging experiments, or optogenetics using bilateral optical cannulas.** Peri-procedural analgesic treatment and anesthesia were carried out the same way as for stereotactic TMEV or AAV injection, described above. An electrode consisting of tungsten wire (Goodfellow Cambridge Ltd., Ø 0.125 mm, 0.014 mm polyimide isolation), crimped with a gold-plated pin (Mill-Max Manufacturing Corp., series 0489), was first cut to a suitable length to be inserted above CA1 (recording electrode), or the cerebellum (reference electrode). Then, the recording electrode was either lowered to CA1 just upon TMEV transduction using the same burr hole established for TMEV injection (coordinates from bregma: AP = -1.9 mm, ML = ±1.6 mm), or via a newly established small burr hole (using a dental drill) above dorsal hippocampus contralateral to the chronic cranial window. Through another burr hole above the cerebellum, a reference electrode of the same type as the recording electrode was inserted. Both electrodes were secured in place using dental cement. In the case of bilateral optical cannula implantation for optogenetic experiments without a cranial window, in addition to the electrodes, optical cannulas (Ø 200 µm, NA 0.5, Thorlabs) were implanted above each dorsal hippocampus, and secured in place using dental cement (Tetric EvoFlow).

**Electrode implantation for chronic wireless EEG recordings.** Peri-procedural analgesia and anesthesia were realized as described above for stereotactic TMEV or AAV injection. Then, mice were implanted bilaterally with depth EEG electrodes in both CA1 regions immediately after TMEV injection, as previously described in detail[16]. Stainless steel screws above the cerebellum (coordinates from bregma: -6 AP, 0 ML, 0 DV) were used as reference electrodes. The correct position of the electrodes was determined histologically in each animal after completion of the respective experiment. Electrographic signals in TMEV-injected animals were recorded using a telemetric EEG monitoring system (Data Science International [DSI], St Paul, MN, USA, sampling rate 1 kHz, bandpass-filtered at 0.5-80 Hz) as described previously[17].

**Implantation of tetrodes and optical cannulas for optogenetic experiments.** To investigate the effect of SD on single unit activity in vivo, adult wildtype bl6 mice received AAV injections to express ChR2-mCherry unilaterally in CA1, as described above. Three weeks after AAV injection, mice were implanted with a modified version of the Versa Drive (Axona) housing one 200 µm light fiber (NA 0.37, Thorlabs), and 4 individually movable tetrodes, each consisting of 4x 0.0127 mm thick tungsten wires (California Fine Wire Company) twisted together[10]. Prior to implantation, tetrodes were gold-plated to an impedance of around 250 kOhm. Mice received an i.p. injection of buprenorphine (0.05 mg / kg body weight) 30 min before surgery start, and were anesthetized with isoflurane (initially 3-4% p.p.a., then reduction to 1-1.4%). The hair above the surgery site was removed, the skin disinfected using iodine solution, and 10 % lidocaine applied as a local anesthetic. To expose the skull, a circular piece of scalp (Ø ~6 mm) was cut out, and all connective tissue was thoroughly removed using 3 % H2O2 and a scalpel. The tissue adhesive Vetbond was applied around the wound and the skull was leveled such that the difference in height between the bone fissures bregma and lambda was less than 0.01 mm. Medio-laterally, the skull was leveled such that the height difference between the centers of the left and right parietal bones (coordinates: AP 2.0, ML +/- 2.0 mm) was less than 0.01 mm. After successful skull alignment, it was covered with the two-component adhesive Optibond and UV-cured for 60 seconds. Next, two craniotomies were established in the interparietal bone (approximate coordinates from bregma: AP 5.5, ML +/- 1.7 mm), and small jeweler screws (00-96X1/16, Protech International Inc) were inserted, serving as anchor screws for the dental cement applied later on. For grounding, two craniotomies were performed in the frontal bone (coordinates from bregma: AP -1.5, ML +/- 2 mm), and the ground screws (jeweler screws, 00-96X1/16, Protech International Inc), soldered to a wire connecting them to the tetrode drive, were inserted with the tip of the shanks facing downwards. Next, a craniotomy of around 1 mm diameter was performed above the right CA1 region using a 0.3 mm drill bit. The underlying dura was carefully removed with a 27G needle, and any potential bleedings were stopped by washing the craniotomy with sterile saline. Once any potential bleeding had stably stopped, the tetrode drive was lowered into the craniotomy such that the light fiber was positioned above CA1 (coordinates from bregma: AP 1.9 ML 1.6 and DV 1.2), and the tetrodes were placed in neocortex. A melted 50:50 paraffin/mineral oil mixture was applied around the tetrodes to prevent the brain from drying, and to avoid that dental cement could come in contact with the brain or the tetrodes. The tetrode drive was fixed to the skull and the anchor screws using air-curing dental cement (Paladur). Postoperative analgesic treatment was carried out the same way as for stereotactic TMEV or AAV injection, described above. Animals typically recovered from surgery within 24–48 hours, showing normal activity and no signs of pain or distress. At least one week of recovery time was allowed before habituation of the mice to the respective experimenters and experimental setup.

**Surgical procedure for 1-photon neocortical widefield imaging and electrophysiology.** Animals were anesthetized and placed in a stereotaxic frame with a nasal mask delivering isoflurane at 0.5–1.5% p.p.a.. During the entire surgery, the body temperature was maintained around 37 °C by a heating-pad placed under the body. For analgesia, animals received a subcutaneous injection of Carprofen (4 mg/kg, Rimadyl, Zoetis GmbH) and Buprenorphine (0.1mg/kg, Buprenovet sine, Bayer Vital GMBH), 20 minutes before the surgical procedure. The hair over the skull was removed and cleaned using phosphate buffered saline (PBS). As an additional local analgesic, Bupivacaine (0.08 ml) was injected under the scalp. The skin over the skull was cut with an incision along the midline, pushed to the sides and attached to the skull using tissue adhesive (Vetbond, 3M). Then, AAV-ChR2 injection into hippocampal CA1 was carried out as described above. For hippocampal electrophysiological recording and optogenetic stimulation, a custom-made optrode, containing a tungsten wire soldered with a gold pin (Goodfellow Cambridge Ltd., 0.125 mm diameter, 0.014 mm polyimide isolation; gold-plated pin: Mill-Max Manufacturing Corp., series 0489), and an optical fiber (Ø 200 µm, NA 0.37) were chronically implanted into the left hemisphere. The tip of the electrode was placed around 200 µm lower than the optical fiber, and the insulation of the tip was removed. Another tungsten electrode soldered with a gold pin was placed on the superficial region of the cerebellum (behind Lambda), as a ground and reference. A light-curing cement was used to secure the implants (Tetric EvoFlow). Dental cement (Superbond [Parkell], Ortho-Jet [Lang Dental]) was then used to build an outer wall for placement of the head bar. A custom-designed head bar was placed parallel to the skull at the level of the dental cement and stabilized with more dental cement around the edges. The skull was then cleared by applying a thin layer of cyanoacrylate (Zap-A-Gap CA+, Pacer Technology)[18]. After the surgery, mice were returned to their home cage on a heating pad, and received the same injections of Carpofen and Buprenorphine as in the beginning for their immediate recovery. They also received Buprenorphine (0.009 mg/ml, Buprenovet sine, Bayer Vital GmbH) and

Enrofloxacin (0.0227 mg/ml, Baytril 5%, Bayer Vital GMBH) in their drinking water for 3 days.

**Animal habituation to 1p- and 2p-microscopes.** Once mice had recovered from surgical procedures for at least two weeks, as evidenced by normal activity and stabilized weight, habituation on a belt (two-photon microscope) or a platform (one-poton widefield microscope) began. Initially, mice were put under the microscope for an interval of up to 30 minutes, freely during the first days, allowing free exploration. Later, they were head-fixed, but still allowed to move freely at will. This habituation was continued until any signs of distress were no longer present. Typically, mice habituated over the course of a few days of training. For two-photon imaging, the individual belt a mouse was initially habituated on was used for all subsequent imaging sessions of that mouse.

**Multimodal two-photon imaging experiments**

**Two-photon $Ca^{2+}$ imaging.** For monitoring neuronal activity in either hippocampus (CA1, stratum pyramidale) or neocortex (layer 2/3, motor cortex), a commercially available two-photon microscope (A1 MP, Nikon) with a tunable laser (Chameleon Vision, Coherent), and a 16x objective (Nikon CFI75 LWD 16X W 0.80 NA 3.0 mm WD) was used. Resonant galvanometer scanning and image acquisition (effective frame rate ~15.2Hz, 512 x 512pixels, ~100µm beneath the hippocampal surface [CA1], or 100-200µm beneath the pial surface [CTX]) were controlled by Nikon imaging software (Nikon NIS-Elements 4.30). Mice with unsatisfactory signal quality after recovery from window implantation were sacrificed according to 3R guidelines, in keeping with European law (2010/63/EU). For viral encephalitis (TMEV) experiments, first, two baseline recordings of 20 minutes each were recorded across 7 days. The field of view was matched across imaging sessions based on stably identified fluorescent neurons, vasculature, FOV location with respect to the bottom of the implanted silicon cone, and depth from hippocampal surface. Immediately after the second baseline imaging, TMEV (or control) injection and electrode insertion were carried out (both contralateral to the imaging window). From day 1-7 p.i., daily multimodal recording sessions (imaging, electrophysiology, locomotion tracking) over up to 3 hours were carried out (several 20min sessions with brief intermittent breaks). After day 7 p.i., recording sessions were reduced to once every 1-2 weeks (experimental duration max. 3 months p.i.). Transgenic Thy1-GCaMP6s animals were imaged at 940-980 nm; jRGECO1a (AAV) mice were imaged at 1060 nm.

**Locomotion tracking on belt.** Locomotion along the belt was tracked with a sensor (Avago Technologies ADNS-9500) detecting the rotation of one of the two belt guide-wheels. The signal was passed through one channel of a 2-channel MiniDigi Digitizer (MiniDigi 1A, Axon Instruments, CA, USA, see LFP methods section below). The recorded rotation signal was then used to calculate % time locomotion, travelled distance, number of locomotion episodes, and maximum speed of locomotion.

**Local field potential (LFP) recordings.** Initially, prior to our first optical detections of seizure-related SD, LFP signals that were recorded alongside two-photon imaging during viral encephalitis, were amplified using a Model 3600 extracellular amplifier (A-M Systems, WA, USA; 0.3 Hz high-pass-, 1 kHz low-pass-filter). Later on, LFP recordings were carried out with a custom-produced EXT-02 B amplifier (Npi Electronic Instruments, Germany; 0.01 Hz high-pass-, 1 kHz low-pass-filter). Regardless of the amplifier used, the differential LFP signal (recording/reference electrode) was digitized at

1kHz by a MiniDigi 1A digitizer (Axon Instruments, CA, USA). The second channel of the digitizer was used for the tracked locomotion signals from the belt. Both channels were recorded using the AxoScope 10.6.2 software (Molecular Devices, CA, USA).

**Multimodal one-photon imaging experiments**

**One-photon widefield $Ca^{2+}$ imaging.** Six to eight weeks post AAV-ChR2 transduction, awake animals were placed into the recording setup for hippocampal LFP recording in combination with neocortical widefield imaging. Widefield imaging was done as previously described[2], using an inverted tandem lens macroscope with the focal lengths of the bottom lens (85M-S, Rokinon) 85 mm and the top lens (DC-Nikkor, Nikon) 105 mm in front of a sCMOS camera (Edge 5.5, PCO). Fluorescence excitation was provided by a collimated blue LED (470 nm, M470L3, Thorlabs) and a collimated violet LED (405 nm, M405L3, Thorlabs) that were coupled into the same excitation path using a dichroic mirror (87-063, Edmund optics). Illumination from the two LEDs was alternated from frame to frame at the frequency 30 frames per second (fps). Using light at two different wavelengths allowed us to isolate $Ca^{2+}$-dependent fluorescence from intrinsic signals (for example, due to hemodynamic responses) by subtracting frames with violet illumination from preceding frames with blue illumination. Fluorescence emission was captured by 525-nm bandpass filter (86-963, Edmund optics) mounted in front of the sCMOS camera. Synchronized to the cortical widefield imaging, LFP signals from hippocampal CA1 were acquired using a 32-channel headstage and a multichannel recording system (ME2100-HS32-M-3m, Multi Channel Systems). The raw signal was then recorded with a passband of 0.01–7500 Hz and a sampling rate of 20 kHz.

**Optogenetic stimulation (with 2p or 1p imaging, or bilateral cannulas).** For one-photon (1p) optogenetic stimulation of hippocampal CA1 in combination with two-photon (2p) imaging, locomotion and LFP recordings (2P/1P/LFP approach), one of two additionally coupled-in CW Chameleon Sapphire LTX lasers was used (488 nm for experimental stimulation, power at brain surface 4-5mW/mm$^2$; or 561 nm for control stimulation, 7 mW/mm$^2$). For the time of 1p-stimulation, the photomultiplier tubes (PMTs) were transiently shut off (thus, 2p-imaging was interrupted for the time of stimulation). Based on published reports[19,20], initial 2P/1P/LFP experiments included titration sessions of varying square-wave 1p-light pulses, to identify suitable and reliable optogenetic ChR2 stimulation parameters to induce Sz (typically 0.5-0.75 sec) or SD (typically 4-4.5 sec). Neither 2p-imaging in the absence of 1p-stimulation, nor control stimulation ever induced Sz or SD. For the optogenetic stimulation of both CA1 regions using bilateral optical cannulas (1p/LFP approach) without 2p-imaging, an LED source was used (Chrolis®/Thorlabs: 475 nm or 590nm [2-3mW/mm$^2$]) using the same stimulation durations for Sz or SD as described above. For optogenetic stimulation of CA1 in combination with neocortical 1p widefield imaging, a multi-wavelength laser module (READYBeam Bio2, FISBA AG) was used to provide light stimulation at either 488 or 635 nm, with a power density of 15 mW/mm$^2$. Stimulation consisted of a continuous, 5-s long light pulse. For all three optogenetic approaches just described, at maximum two optogenetic stimulations (1 control stimulation followed by 1 experimental stimulation) were performed per day (in total 3-6 stimulations per condition [ctr, Sz, SD] per mouse). Optogenetic experiments were always started once mice were completely relaxed on the belt/platform (~30min after being transferred). Individual multimodal recordings sessions always spanned 15 min (stimulation after 5 min baseline).

**Tetrode recordings and optogenetic stimulation.** 7 days after drive implant surgeries, mice were habituated to the experimenter and to be restrained at the tetrode drive. After successful habituation, mice were ready for recordings. The recording setup consisted of a Neuralynx recording system (Digital Lynx 4SX) that was connected to the tetrode drive via a 16-channel head-mounted amplifier and a 4 m cable. The recording system was further connected to a computer running the recording software Cheetah 5.0, and the behavior tracking software Noldus 14. LFP was recorded at 32000 Hz, lowpass bandpass filtered between 0.1 and 800 Hz, and amplified 1000x. For spike detection, signals were band-pass filtered at 600 and 6000 Hz and waveforms were time-stamped if they exceeded 40 µV.

For optogenetic SD stimulation, a TTL pulse was sent via the Noldus system to a digital acquisition system (Molecular Devises) that in turn sent a 3.5-second-long pulse to a 100 W 473 nm laser (Omicron-Laserage). The laser was connected to the light fiber protruding the tetrode drive via a patch cord (NA 0.39, Thorlabs). The TTL pulse was split such that it additionally sent a timestamp to the Neuralynx system. The laser power was set such that the output of the implanted light fiber was ~1.6 $mW/mm^2$.

To record single unit activity from CA1 neurons, one reference tetrode remained in cortex while the other tetrodes were slowly lowered (~50 µm/day) until theta oscillations, sharp-wave-ripples and spikes were detected, indicating that CA1 was reached. Each recording consisted of a 20 min baseline recording, followed by a 3.5 second optogenetic stimulation to initiate SD, followed by another 20 minutes of post-stimulation recording. One recording session was performed per animal per day, and recordings were repeated over multiple days. When spiking activity disappeared between recordings sessions, or the spike amplitude became too low, tetrodes were turned slightly deeper until new spikes appeared. During the recordings, mice were placed in a circular glass chamber.

**Immunohistochemistry (IHC) and image acquisition.** For IHC, mice were deeply anesthetized with 16mg/kg xylazine (20mg/mL, i.p., xylazine hydrochlorid, Serumwerk Bernburg AG) and 100mg/kg ketamine (10%, Serumwerk Bernburg AG), and transcardially perfused with PBS and 4% formaldehyde (FA, in PBS). Brains were removed and stored overnight in 4% FA at 4°C. Coronal sections (40 µm) were used for immunofluorescent stainings against Iba1 (FUJIFILMS Wako Chemicals, 019-19741, 1:200), GFAP (Abcam, ab53554, 1:200), and CD8 (Abcam, ab10558, 1:100).

Briefly, slides were washed twice in PBS and once in PBS containing 0.25% Triton (PBT) for 10 min, subsequently blocked for 1 hour at RT with PBT containing 10% normal goat (NGS) and donkey (NDS) serum, respectively, depending on the secondary antibody, and then incubated with primary antibodies diluted in 3% serum/PBS overnight at 4°C. After washing three times with PBT for 10 min, slides were incubated with respective secondary antibodies (AlexaFluor555-conjugated donkey anti-goat, Invitrogen, A32816; AlexaFluor647-conjugated goat anti-rabbit, Invitrogen, A21247; AlexaFluor647-conjugated goat anti-rabbit, Life Technologies, 1445259) for 1.5 h at RT in 3% serum/PBS. For counter-staining, slides were then washed twice with PBT for 10 min at RT, incubated with DAPI (4', 6-Diamidino-2-phenylindole dihydrochloride, ThermoFisher Scientific, 1:1000 in PBS) for 5 min at RT, and mounted with Aqua-Poly/Mount (Polysciences). The confocal microscope Visiscope (Visitron Systems GmbH) attached to the spinning disc unit CSU-W1 (Yokogawa) was used for image acquisition. Images were digitized with pco.edge sCMOS cameras (Excelitas

Technologies), and visualized with the VisiView®-Software (Visitron Systems GmbH). Acquired images were further analyzed using FIJI (ImageJ).

#### Analysis of human depth-electrode and microwire recordings

##### Electrographic signal recordings

Behnke-Fried (BF) depth electrodes were implanted following a pre-surgical plan based solely on clinical considerations, with the most distal macro-contact placed closest to target regions of interest. BF microwire bundles (8 recording wires per bundle, one bundle per depth electrode) were implanted in addition to clinical depth electrodes (each carrying 8 macro-contacts across depth). BF microwires can be targeted to regions of interest through the hollow core of clinical depth electrodes (Ad-Tech®, WI, USA), and have no additional bearing on the risk of adverse effects or complications related to depth electrode placement itself[21].

Microwire bundles protruded around 4 mm from the electrode's tip. Data were recorded using a Neuralynx ATLAS amplifier and Pegasus software (Neuralynx, Bozeman, MT, USA) with a 0.1 Hz – 9 kHz band-pass filter, sampled at 2 kHz for macro-contacts and at 32 kHz for microwires, and stored locally. Further analysis was carried in 60-minute-long cuts of data centered at the earliest regional seizure onset occurrence. Macro-contact traces were re-referenced to a bipolar reference scheme. From microwire recordings, potential spikes were extracted and clustered into putative Units using the Combinato software[22] and later visually confirmed and separated into Multi-Units (MU) and Single-Units (SU). Every cluster's mean waveform was removed from each of its spike occurrences in the recorded microwire signal. Local Field Potential traces (LFP) were obtained by re-sampling the mean-spike-removed microwire signal to 1 kHz, low-pass filtering it (1st order Butterworth low-pass filter, cutoff frequency: 300 Hz) and re-

referencing them to a local common average reference scheme using in-house functions built in Python 3.8.

### Data analysis

#### Animal experiments

Data from animal experiments were analyzed with Python 3.8.12 (code available upon request). Legacy Matlab code (adapted for Matlab R2023a, MathWorks) was used to match the locomotion and two-photon imaging recordings. For statistical analysis, Prism 10.3 (GraphPad) was used. Parametric tests were used if groups were normally-distributed (Kolmogorov-Smirnov normality test, alpha=.05), and non-parametric tests if not.

**Neuronal $Ca^{2+}$ signal extraction.** To obtain the $Ca^{2+}$ signal of single neurons, CaImAn (1.9.7) was used. Due to the excessive non-physiological $Ca^{2+}$ transients during Sz and SD, the standard pipeline had to be modified as follows. First, motion correction was performed over the whole respective recording, and the results were saved. Next, the segments mentioned above were cut out. The rest of the frames were concatenated, and the spatial and temporal components were extracted. The spatial components were used as masks over the whole (motion-corrected) recording: for each component, the value at each pixel (coordinate) was used as a weight to the fluorescence signal of that location. The signal corresponding to the given spatial component was taken as the weighted average signal of the pixels constituting the component.

**Sz and SD onset times at single cell level.** For each spatial component, the center of mass was found and defined as neuron location. Each trace was smoothed using LOWESS-filtering (2s time windows). For each neuron $n$ and each frame $i$, the central finite difference approximation of the first derivative was calculated from the filtered time series ($y_i^n$) as

$$(y')_i^n = \frac{-1}{60} \cdot (y_{i+3}^n - 9y_{i+2}^n + 45y_{i+1}^n - 45y_{i-1}^n + 9y_{i-2}^n - y_{i-3}^n)$$

To achieve more resistance to flashing activity occurring simultaneously with the slowly changing SD signal, an asymmetric difference signal was also calculated:

$$a_i^n = y_{i+4}^n + y_{i+2}^n + y_{i+1}^n - y_{i-1}^n - y_{i-2}^n - y_{i-3}^n$$

Where applicable, a seizure detection window was defined as a window between the beginning and end of observable epileptic activity, or the onset of an SD wave. An SD detection window (SD depolarization part) was defined as starting with the first frame of SD appearance in the imaged field of view, until the entire field of view was optically invaded by SD, and thus, maximum population average fluorescence was reached. The seizure onset frame was found for each cell as the first maximum of $(y')^n$ within the seizure time window. For multiplexed SDs, the number of SD waves $n_{SD}$was visually determined, and the onset frame of each SD wave at cell $n$ was found as the first $n_{SD}$ maxima of $a_i^n$ within the SD detection window (in the overwhelming majority of cases, $n_{SD} = 2$ [sSD1, sSD2]). Based on observations, a minimum temporal distance of 2-5s was also required between two onset time points for a given cell.

**Sz and SD directionality analysis.** For each individual event (Sz, sSD1, sSD2), the 5% largest deviations from the median onset time were discarded in the dataset of cell-level onset time, obtained as described above. Of the remaining onset times, using the corresponding cell locations, the geometrical center $\vec{x}_{ce}$ of the earliest 25% were calculated, and $\vec{x}_{cl}$ was found similarly for the latest 25% onset times. The direction of the event was then defined as

$$\vec{x}_{ed} = \frac{\vec{x}_{cl} - \vec{x}_{ce}}{|\vec{x}_{cl} - \vec{x}_{ce}|}.$$

The mean direction of specific events (*e.g.* the mean direction of seizures $\vec{x}_{ed}^{i}$ for a single mouse) was defined as the direction of the vector sum $\vec{x}_{md} = \sum_{i=1}^{n_e} \vec{x}_{ed}^{i}$, i.e. $\frac{\vec{x}_{md}}{|\vec{x}_{md}|}$. For each mouse *i* and for each event type *j* (Sz, sSD1, sSD2), the standard deviation $\sigma_{i,j}$ of all $n_{i,j}$ measured directional angles was calculated (with care to avoid the wrap-around effect) as a measure of clustering. Each obtained value was compared to a surrogate dataset $S_{i,j}$ of 1000 standard deviations calculated from $n_{i,j}$ uniformly distributed random directional angles. With this measure of clustering, a test of the null hypothesis $H_0$ that the experimentally recorded directions for a given mouse and given event type are uniformly distributed was performed: for each *i* and j, the threshold $\hat{\sigma}_{i,j}$ of 5% lowest standard deviations of $S_{i,j}$ was determined. At the α=0.05 significance level, $H_0$ could be rejected if $\sigma_{i,j} \leq \hat{\sigma}_{i,j}$.

**Sz and SD progression speed.** The progression speed of the SD wave front was estimated using the cell-level onset data. For each neuron $\vec{x}^{n}$ with onset time $t^{n}$, finding the closest neighboring cell $\vec{x}^{c}$ that had a later onset time, that is, $t_c > t_n$, was attempted. If such an $\vec{x}^{c}$ was found, the average speed

$$v^{n} = \frac{1}{t_c - t_n} |\vec{x}^{c} - \vec{x}^{n}|$$

was calculated. Using imaging frequency and microscope resolution data, $v^{n}$ was converted to units of mm/min. The median of the $v^{n}$ values for a given SD event is the estimate of the wave front speed. The onset speed of the seizures was approximated similarly.

**Sz and SD signal amplitudes.** Using the spatially averaged fluorescence signal of the whole field of view, a baseline fluorescence was determined as the median of the lowest 5% values within a 10s window directly before visually confirmed Sz invasion for CA1 recordings (where necessary, manually shifted to a segment without flashing activity), and ~1 minute before visually confirmed Sz onset in neocortical recordings, where invasion occurred in a more fractured way than in CA1. The amplitude of an optical seizure was determined as the mean of the highest 5% fluorescence values within the window of visual Sz detection and appearance of an SD depolarization wave, or the end of seizure activity (if no SD occurred). SD amplitude ($y_{SDamp}$) was calculated based on the time point of absolute maximum fluorescence upon SD appearance in the imaged field of view ($t_{SDamp}$).

**Sz and SD post-ictal recovery time characterization** Starting from the time point $t_{SDamp}$, or, in absence of SD, a manually defined time point ~5-15 s before the apparent depression of $Ca^{2+}$ activity, the subsequent trough of fluorescence was located as the median time point of the 15 earliest of the 5% lowest fluorescence values within 5000 frames (~5.5 minutes) window. The corresponding trough signal amplitude was derived as the median of the lowest 5% values within a 10s window centered around this trough time point. Recordings, where this amplitude was greater than the same metric for the baseline (as described above) were discarded from recovery analysis. To characterize the recovery, amplitudes were calculated in a similar manner for subsequent 10s windows, in 5s shifts of the center, until a window with corresponding amplitude reached 95% of the baseline amplitude of fluorescence. If the end of the recording was reached without finding such a window, the trough and the last window were used for linear extrapolation. If this also failed, the recording was excluded from this analysis. The center point of the first window with the amplitude above 95% of baseline or the extrapolated value were taken as the fluorescence recovery time point (as a proxy for basic neuronal network recovery).

**Half width at half maximum (HWHM) of SD signal.** For comparison of optostimulation-induced and encephalitis-related SD characteristics (see Suppl. Fig. 5), the HWHM on the trailing edge was calculated for each waveform. The SD amplitude ($y_{SDamp}$, corresponding time $t_{SDamp}$) and subsequent trough amplitude ($y_{min}$) were used, both as described earlier. The time point where the signal subsequently reaches $(y_{max} + y_{min})/2$ was determined ($t_{half} > t_{SDamp}$). The HWHM equals $t_{HWHM} = t_{half} - t_{SDamp}$.

**Locomotion analysis.** For analysis of locomotion, two equally-sized windows were always compared. For all imaged or electrically recorded Sz (encephalitis, or optogenetics), a “pre-ictal” baseline window of ~5min (directly preceding Sz) was compared to an equally sized “post-ictal” window (starting at the end of Sz, regardless of Sz with or without SD). For optogenetically stimulated SD and control, pre- and post-ictal time windows of equal size, as just described, were determined based on the start and stop of optogenetic stimulation.

The calculated metrics were proportion of time spent (in %), total distance travelled (arbitrary units, a.u.), number of locomotion episodes, and maximum speed (a.u.). Locomotion episodes were identified by binary labeling each time point as either resting (velocity = 0) or locomotion. Subsequent frames labeled as locomotion formed episodes (an episode may contain a single sample). Episodes separated by less than 0.5

s were merged. In these cases, the beginning of the first merged episode and the end of the last one mark the beginning and end of the resulting episode, and all frames within the segment were classified as frame with locomotion. The resulting new set of episodes is then filtered by two criteria: firstly, during one episode, the velocity must at one point reach a threshold value, which was specified by observing locomotion amplitude during non-locomotive behavior resulting in belt movement (sitting, grooming), and setting the threshold to discard these segments. Secondly, the episode must last longer than 1s.

**Analysis of widefield imaging data.** Widefield data analysis was performed as previously described[18,23,24]. Briefly, data were corrected for motion artifacts of the skull relative to the objective by aligning each frame to a time-averaged reference image. The next step was to compute relative fluorescence changes (ΔF/F) for each pixel to measure small changes in fluorescence due to cortical activity. Computing ΔF/F was done by subtracting and dividing the value at each frame by the baseline fluorescence. We then used singular value decomposition (SVD) to transform imaging data from pixels to data components, reducing the data size and computational cost of further analysis[3]. All subsequent analysis across time was performed on the 500 components that described the highest-variance in the imaging data. For hemodynamic correction, the $Ca^{2+}$-independent fluorescence signals acquired under violet illumination were rescaled and subtracted from the $Ca^{2+}$-dependent fluorescence, according to the approach presented by Allen et al.[25]. The resulting low-dimensional and hemodynamic-corrected signal was used for subsequent analysis. Imaging data were then aligned to the Allen Mouse Brain Common Coordinate Framework (CCF), using anatomical landmarks: the left, center and right points where the cortex contacts the olfactory bulbs and the medial point on the midline at the base of the retrosplenial cortex. The accuracy of CCF alignment was also functionally confirmed through retinotopic visual mapping[23].

To isolate cortical responses, relative to CA1 stimulation, we extracted the stimulus-aligned fluorescence signals from the right hemisphere or specific areas of interest. To quantify changes directly after CA1 stimulation, we compared the median fluorescence for 10 s after stimulus offset with a 10 s baseline window. To identify a potential reduction in fluorescence in different areas, we used a 10-s moving average to remove transients in activity and then computed the time and amplitude of the lowest 5% fluorescence in a 90 s post-stimulus time period. The same procedure was used for a 90 s baseline period. Changes in fluorescence were then computed as the difference between the lowest fluorescence in the baseline minus the postictal period. In case of a reduction in cortical activity after CA1 stimulation, the recovery time was computed as the time until the cortical activity was above the lowest 5% fluorescence in the baseline period (Suppl. Fig. 8).

**Analysis of tetrode recordings.** Units were manually clustered using MClust 3.5 (D. Redish https://redishlab.umn.edu/mclust) in Matlab 2023b (Mathworks) based on the parameters' peak size, energy of spike waveforms and peak-to-valley amplitude. The time stamps of the extracted units as well as the LFP trace of one electrode of the corresponding tetrode were then loaded into Matlab 2023b and further processed using custom-written scripts.

### Analysis of human recordings

#### Identification of Sz-sSD

Sz onset and termination times in the LFP traces were visually adjusted for each region assisted by aligned spectrograms. For each microwire, a period of interest of 30 seconds of its post-ictal LFP trace was low-pass filtered (1st order Butterworth low-pass filter, cut-off frequency: 1 Hz) and mean-corrected. Potential slow-shift components were identified by the following inclusion criteria: the rising phase of the sSD occurred within the first 5 seconds after Sz termination and the absolute amplitude of the sSD exceeded at least half the regional maximum amplitude. Rising phase time was calculated by taking the first derivative's maximum absolute value of the filtered trace. A region was considered sSD-invaded if at least 3 of its 8 microwires showed sSD. All potential sSD invasions were visually confirmed by two experimenters, assisted by unit firing and aligned spectrograms, and cases where regions showed no noticeable decrease in power during the sSD period were removed from the sSD-invaded category.

**Power spectral analysis**

Power spectral analysis of periods of interest was carried out by choosing a 1-minute-long segment preictally (with an added 30 second buffer before adjusted regional LFP Sz onset) and postictally (for sSD-invaded traces, the start of this period was adjusted to avoid the duration of the sSD itself). The spectral content of the segments was obtained by calculating consecutive Fourier transforms using scipy's spectrogram function (Python 3.8, scipy 1.10.1 signal toolbox). A baseline period for normalization of the spectral data was chosen to be the average of 10 non-overlapping 1 minute-long randomly chosen periods, occurring between 30 and 10 minutes prior to each Sz. Normalisation was done as follows:

$$PSD_{norm} = 10 * log_{10}(PSD_{period}/PSD_{baseline})$$

where PSDperiod and PSDbaseline are the power spectral density of the period of interest and the baseline, respectively, and PSDnorm is the normalized power spectral density, in decibels (dB). The energy content of each period was calculated as the average normalized decibels between 2-50 Hz (broadband) and 80-150 Hz (high gamma)[26]. Additionally, the same energy value was calculated for 15 1-minute-long periods postictally for each of the power bands, and a linear regression was performed in order to calculate a time of recovery to its average preictal value. Regional values were calculated as an average of the power across all channels within the region. Comparisons between Sz-invaded and Sz-sSD-invaded regions were carried out for a pool of seizures where at least one region experienced sSD using a paired t-test when both groups were normally-distributed (Kolmogorov-Smirnov normality test, alpha=.05) and a Mann-Whitney U test if not. Statistics were run using Prism 10.3 (GraphPad) and Python's scipy 1.10.1 stats toolbox.

**Materials and Methods Section – Additional References**


1. Dana, H. *et al.* Thy1-GCaMP6 transgenic mice for neuronal population imaging in vivo. *PLoS One* **9**, e108697 (2014).

2. Masala, N. *et al.* Aberrant hippocampal Ca2+ micro-waves following synapsin-dependent adeno-associated viral expression of Ca2+ indicators. *Elife* (2024). doi:https://doi.org/10.7554/eLife.93804.1

3. Wekselblatt, J. B., Flister, E. D., Piscopo, D. M. & Niell, C. M. Large-scale imaging of cortical dynamics during sensory perception and behavior. *J. Neurophysiol.* **115**, 2852–2866 (2016).

4. Libbey, J. E. *et al.* Seizures following picornavirus infection. *Epilepsia* **49**,

1066–1074 (2008).
