## Supplementary material for "Region-specific spreading depolarization drives aberrant post-ictal behavior": Suppl_Data_Legends

### Suppl. Figures, Tables and Movie Captions

**Suppl. Fig. 1. Histology of TMEV encephalitis and subsequent histological features of temporal lobe epilepsy**

**A**, DAPI (blue) and Iba-1 (red) staining. Top: control 3 months post mock injection. White arrowhead indicates injection site. Chronic cranial hippocampal window for chronic 2p imaging of CA1 in contralateral hemisphere. Middle: 3 days post injection of TMEV (arrowhead). Bottom: 14 days post injection of TMEV. **B**, magnified insets from stained slices in A. Note enlarged hyper-ramified microglia during early encephalitis (d3), and severe inflammation with amyboid microglia during late encephalitis (d14). **C**, DAPI (blue) and CD45 (red) staining. Top: control 3 months post mock injection. White arrowhead indicates injection site. Chronic cranial hippocampal window for chronic 2p imaging of CA1 in contralateral hemisphere. Middle: 3 days post injection of TMEV (arrowhead). Bottom: 14 days post injection of TMEV. **D**, magnified insets from stained slices in C. Note increasing presence of CD45 cells in hippocampus across encephalitis, in a delayed fashion in comparison to A, B. **E**, DAPI (blue) and GFAP (green) staining. Top: control, 3 months post mock injection. White arrowhead indicates injection site. Chronic cranial hippocampal window for chronic 2p imaging of CA1 in contralateral hemisphere. Bottom: 3 months post-TMEV injection. Chronic cranial hippocampal window for chronic 2p imaging of CA1 in contralateral hemisphere. Note the massive astrogliosis and severe hippocampal tissue damage, in line with known hallmarks of post-encephalitic temporal lobe epilepsy in humans. **F**, magnified insets from stainings in E.

**Suppl. Fig 2. Bi-hippocampal wireless EEG recordings during TMEV encephalitis**

Waterfall plot of respective 2 min windows across 29 visually identified electrographic Sz across 5 days during encephalitis. Blue = left hippocampal electrode, black = right hippocampal electrode. Note that all electrographic Sz are detected bilaterally. EEG recording: 0.5 Hz HP-, 80 Hz LP-filter.

**Suppl. Fig 3. Sz- versus sSD-related locomotion during encephalitis**

Comparison of locomotory parameters during Sz vs. following sSD in hippocampal imaging during viral encephalitis. **A**, Locomotion recording (belt) and 2P-imaging of 3 encephalitic hippocampal Sz-sSD in an example mouse. To compare locomotion in equally sized temporal windows across optical Sz and sSD, window lengths for both Sz and sSD were determined and matched based on the shorter event duration (Sz durations consistently shorter than sSD). Note that while Sz-duration-matched window sizes were required for proper statistical comparison, the biological duration of sSD (minutes) outlasted Sz-duration by far. **B-D**: Comparison of 11 Sz- vs. 11 sSD periods in all CA1-imaged 4 mice (two events excluded from analysis due to brief intermittent imaging break during Sz, see Fig. 1 F left). **B**, % time locomotion, paired t-test (1.515 ± 1.515 [Sz] vs. 13.74 ± 5.676 [sSD], p=0.0259). **C**, Travelled distance (a.u.: arbitrary units), paired t-test (0.078 ± 0.044 [Sz] vs. 0.567 ± 0.223 [sSD], p=0.0459). **D**, maximum speed (a.u.), paired t-test (0.107 ± 0.038 [Sz] vs. 0.281 ± 0.069 [sSD], p=0.0427). For entire Fig.: All given ± denote s.e.m., all bar plots represent means ± denote s.e.m.. Depiction of statistical significance: *p < 0.05, **p < 0.01.

**Suppl. Fig. 4. Optogenetic SD induction and 2P imaging using jRGECO and ChR2**

**A**, Optogenetic experimental setup: 2p/1p/LFP approach, CA1 window for 2p neuronal $Ca^{2+}$ imaging (jRGECO1a) and 1p optogenetic stimulation (ChR2-GFP), both contralateral to LFP electrode (black pin, at CA1). **B**, Co-expression of ChR2 (AAV, green) and jRGECO (AAV, magenta) in CA1 (str. pyr.). **C**, Example of optogenetic SD induction: 4 sec optogenetic stimulation (488 nm) triggers isolated SD in imaged field of view. In contralat. LFP, neither Sz nor SD are detected. Top trace depicts avg pop. $Ca^{2+}$ signal, middle traces LFP (0.01 HP filter), bottom trace locomotion.

**Suppl. Fig 5. SD features in TMEV vs. optogenetic experiments**
**A**, Comparison of sSD/Sz amplitude ratio in TMEV encephalitis and optogenetic stimulation (ChR2). Unpaired t-test (12 Sz-sSD events in TMEV vs. 9 optogenetic Sz-sSD events, 1.453 ± 0.036 vs. 1.43 ± 0.281, p=0.9269). **B**, Comparison of optogenetic sSD and optogenetic SD depolarization amplitude half-width. Mann-Whitney-test (9 sSD vs. 16 SD, 10.22 ± 1.56 vs. 15.4 ± 1.749, p=0.357). **C**, Comparison of optogenetic sSD and optogenetic SD speed (SD depolarization wave). Mann-Whitney-test (6 sSD [3 events excluded due to complex spatiotemporal expansion] vs. 8 SD [8 events excluded due to missed optical onset during stim.], 4.954 ± 0.1868 vs. 5.835 ± 1.357, p=0.4908). **D**, CA1 tetrode recordings (3 mice [gray shades], 17 optogenetic SD) of LFP (top half, and corresponding avg spectrogram) and corresponding single unit activity (SUA) (lower half, same mice [gray shades], vertical black lines show recorded single units per SD stimulation). Blue shade depicts optogenetic stimulation (473nm, ~5 $mW/mm^2$, 3.5 sec square wave illumination). Magnified inset of LFP SD stimulations on the right (green dotted box). Note the cessation of firing of most single units upon SD, followed by sustained depression of firing.
For entire Fig.: All given ± denote s.e.m.. Depiction of box plots: mean (solid line), boxes extend from $25^{th}$ - $75^{th}$ percentiles. Depiction of statistical significance: n.s. not significant, *p < 0.05, **p < 0.01, ***p < 0.001, ****p < 0.0001.

**Suppl. Fig 6. Locomotory phenotype in two different optogenetic Sz/SD experimental runs**
**A-D**, 2P-1P-LFP approach [see figure 2 G, left], 4 mice (12 control [ctr], 17 SD, 9 Sz-sSD), exclusively unilateral (uni.) stimulations, max. one SD and Sz-sSD stim. per day. **A**, Δ % time locomotion, -6,322 ± 1.158 (ctr), 8.331 ± 1.861 (SD), 38.11 ± 4.861 (Sz-sSD). One-way ANOVA with Tukey's test (F[2, 35]=64.11; ctr vs SD p=0.0003, ctr vs Sz-sSD p<0.0001, SD vs. Sz-sSD p<0.0001). **B**, Δ travelled distance (a.u.): -2.810 ± 0.6701 (ctr), 5.027 ± 1.063 (SD), 27.66 ± 4.549 (Sz-sSD). One-way ANOVA with Tukey's test (F[2, 35]=47.42; ctr vs SD p=0.0193, ctr vs Sz-sSD p<0.0001, SD vs. Sz-sSD p<0.0001). **C**, Δ # of locomotion episodes: -3.25 ± 0.653 (ctr), 4.176 ± 1.129 (SD), 10.0 ± 1.155 (Sz-sSD). One-way ANOVA with Tukey's test (F[2, 35]=32.69; ctr vs SD p<0.0001, ctr vs Sz-sSD p<0.0001, SD vs. Sz-sSD p=0.0018). **D**, Δ maximum speed (a.u.): -0.238 ± 0.0686 (ctr), 0.072 ± 0.063 (unilat. SD), 0.257 ± 0.087 (Sz-sSD). One-way ANOVA with Tukey's test (F[2, 35]=10.52; ctr vs SD p=0.007, ctr vs Sz-sSD p=0.0003, SD vs. Sz-sSD p=0.1959. **E-H**, 1P-LFP approach [see figure 2 G, right], 3 mice (11 uni. ctr, 10 bilat. ctr, 7 unilat. Sz-sSD, 4 bilat. Sz-sSD, 10 bilat. SD), unilateral (uni.) or bilateral (bilat.) stimulations, max. one SD and Sz-sSD stim. per day; comb. (combined) denotes pooled data (from groups marked in blue boxes, respectively). Statistics are given for pooled data (for unpooled data, see Suppl. Table 1, for reasons of practicality). **E**, Δ % locomotion, -3.306 ± 1.72 (ctr), 27.33 ± 4.755 (Sz-sSD), 40.68 ± 4.325 (bilat. SD). One-way ANOVA with Tukey's test (F[2, 39]=55.13; ctr vs Sz-sSD p<0.0001, ctr vs. bilat. SD p<0.0001, Sz-sSD vs. bilat. SD p=0.0348) **F**, Δ travelled distance (a.u.): -1.231 ± 0.8318 (ctr), 18.82 ± 2.893 (Sz-sSD), 25.95 ± 3.159 (bilat. SD). One-way ANOVA with Tukey's test (F[2, 39]=55.83; ctr vs. Sz-sSD p<0.0001, ctr vs. bilat. SD p<0.0001, Sz-sSD vs. bilat. SD p=0.0808. **G**, Δ # of locomotion episodes: -

0.8095 ± 0.5461 (ctr), 6.455 ± 1.296 (Sz-sSD), 7.0 ± 1.461 (bilat. SD). One-way ANOVA with Tukey's test (F[2, 39]=23.15; ctr vs. Sz-sSD p<0.0001, ctr vs. bilat. SD p<0.0001, Sz-sSD vs. bilat. SD p=0.9355. **H**, Δ maximum speed (a.u.): -0.102 ± 0.071 (ctr), 0.230 ± 0.125 (Sz-sSD), 0.366 ± 0.106 (bilat. SD). One-way ANOVA with Tukey's test (F[2, 39]=7.067; ctr vs. Sz-sSD p=0.039, ctr vs. bilat. SD p=0.0037, Sz-sSD vs. bilat. SD p=0.6609. **I-L**, Combined 2P-1P-LFP ($1^{st}$ set of exp., black) and 1P-LFP ($2^{nd}$ set of exp., pooled data, magenta) exp. shown in A-H; in total 7 mice (12 ctr [set 1], 21 ctr [set 2], 9 Sz-sSD [set 1], 11 Sz-sSD [set 2], 17 unilat. SD [set 1], 10 bilat. SD [set 2]). One-way ANOVA with Šidák's multiple comparisons test, for detailed statistical results please be referred to Suppl. Table 2, for reasons of practicality. Note: The resulting final data from pooling of non-significantly different control (uni- vs. bilateral stim.) or Sz-sSD (uni- vs. bilateral stim.) conditions in suppl. Fig. 6 I-L are shown in Fig. 2 K-N. For entire Fig.: All given ± denote s.e.m.. Depiction of violin plots: median (solid lines), quartiles (dotted lines). Depiction of statistical significance: n.s. not significant, *p < 0.05, **p < 0.01, ***p < 0.001, ****p < 0.0001.

**Suppl. Fig 7. Raw locomotory traces across all mice in two different optogenetic Sz/SD experimental runs**
Raw locomotory data from 2P-1P-LFP and 1P-LFP stimulation experiments, combined (7 mice [gray shades]; 33 control, 17 SD unilateral, 20 Sz-sSD, 10 SD bilateral. for statistical comparisons, see Suppl. Fig. 6, and Suppl. Tables 1 and 2.

**Suppl. Fig 8. Calculation of post-stim. signal trough and recovery in 1p widefield imaging**
Paradigmatic example. Top: Hippocampal (HPC) LFP of optogenetic SD stimulation (488nm, 5 sec, blue shade), and cortical widefield $Ca^{2+}$ activity (Δ F/F) in secondary/primary motor (M2, M1) and parietal cortex (P). Gray traces show the smoothed (30sec moving avg) $Ca^{2+}$ signal that was used to calculate a post-stim minimal value (red dotted lines) and the trough to recovery time to pre-stim values (blue dotted line).

**Suppl. Fig 9. The clinical EEG filter standard renders SD invisible**
*In vivo* LFP recording of one optogenetically induced Sz-sSD in mouse CA1. The same electrographic Sz-sSD is passed through different high-pass filters (0.01 - 0.5 Hz). Note that at ≥0.3 Hz, sSD is no longer visible in the signal trace (EEG standard ≥0.5 Hz).

**Suppl. Fig 10. Reconstructed brain MRI and location of implanted electrodes, and clinical data**
**Top**, 3D-reconstructed brain MRI and location of implanted BF electrodes. Green dots indicate macro-contact locations, shades of green designate different brain regions and violet markers depict microwire bundles., AMD: amygdala, aHPC: anterior hippocampus, EC: entorhinal cortex, pHPC: posterior hippocampus, PHC: parahippocampal cortex, PIC: piriform cortex, Ta: temporal cortex a, Tb: temporal cortex b, Tc: temporal cortex c. **Bottom**, Corresponding to the 3-D reconstructions, clinical data for each patient included in the study. FCD: focal cortical dysplasia.

**Suppl. Table 1. Statistical comparison of sub-groups from unpooled data shown in suppl. Fig. 6 E-H**

**Suppl. Table 2. Statistical comparison of sub-groups from data shown in suppl. Fig. 6 I-L**

**Suppl. movie 1. Two-photon imaging of CA1 Sz invasion during encephalitis**
Two-photon calcium imaging in a transgenic thy1-GCaMP6s mouse in hippocampal CA1, around 100 µm beneath hippocampal surface (stratum pyramidale), on day 2 of viral encephalitis (note: TMEV transduction in contralateral CA1, see Fig. 1A) FOV ~700x700 µm, Imaging wavelength = 940 nm, acquisition speed = 15 frames/sec. Movie played at 2x acquisition speed. Note, Sz is followed by sSD.

**Suppl. movie 2. Two-photon imaging of CTX Sz invasion during encephalitis**
Two-photon calcium imaging in a transgenic thy1-GCaMP6s mouse in neocortex (motor region), around 150 µm beneath pial surface (L II/III), on day 2 of viral encephalitis (note: TMEV transduction in contralateral CA1, see Fig. 1A) FOV ~700x700 µm, Imaging wavelength = 940 nm, acquisition speed = 15 frames/sec. Movie played at 2x acquisition speed. Note the absence of sSD post Sz.

**Suppl. movie 3. Optogenetic Sz-sSD induction in CA1 (2p/1p/LFP approach)**
Two-photon calcium imaging in a transgenic thy1-GCaMP6s mouse in hippocampal CA1 (around 100 µm beneath hippocampal surface [stratum pyramidale], FOV ~700x700 µm, Imaging wavelength = 940 nm, acquisition speed = 15 frames/sec.), with additional AAV-hSyn-hChR2[H134R]-mCherry, Addgene ID 26976-AAV5 in CA1, around six weeks post transduction (see Fig. 2 G). Optogenetic induction of Sz-sSD via brief square wave light pulse (750 ms, 488 nm, 4-5mW/mm$^2$, see fig. 2H, left). Movie played at 2x acquisition speed. Note, Sz is followed by sSD.

**Suppl. movie 4. Optogenetic SD induction in CA1 (2p/1p/LFP approach)**
Two-photon calcium imaging in a transgenic thy1-GCaMP6s mouse in hippocampal CA1 (around 100 µm beneath hippocampal surface [stratum pyramidale], FOV ~700x700 µm, Imaging wavelength = 940 nm, acquisition speed = 15 frames/sec.), with additional AAV-hSyn-hChR2[H134R]-mCherry, Addgene ID 26976 in CA1, around six weeks post transduction (see Fig. 2 G). Optogenetic induction of SD via square wave light pulse (4000 ms, 488 nm, 4-5mW/mm$^2$). Movie played at 2x acquisition speed. Note isolated SD induction, no Sz occurrence.

**Suppl. movie 5. Optogenetic SD induction in CA1 (2p/1p/LFP approach)**
Two-photon calcium imaging in hippocampal CA1 (around 100 µm beneath hippocampal surface [stratum pyramidale], FOV ~700x700 µm, Imaging wavelength = 940 nm, acquisition speed = 15 frames/sec.), around 6 weeks post AAV-transduction of jrGECO1a and ChR2 in CA1 of a bl6 wildtype mouse (AAV2/1-CAMKIIa-jNES-jRGECO [Uni Bonn Vector Core] and AAV-Syn-hChR(H134R)-GFP [Addgene_58880], see Suppl. Fig. 4). Optogenetic induction of SD via square wave light pulse (4000 ms, 488 nm, 4-5mW/mm$^2$). Movie played at 2x acquisition speed. Note isolated SD induction, no Sz occurrence.

**Suppl. movie 6. Optogenetic SD induction in CA1 and 1p widefield CTX imaging**
One-photon imaging of the whole dorsal cortex through cleared skull in a transgenic CaMK2α-tetO-GCaMP6s mouse (FOV = 12.5x10.5 mm$^2$, Imaging wavelength = 470 nm, acquisition speed = 15 frames/sec.) with additional AAV-hSyn-hChR2[H134R]-mCherry, Addgene ID 26976 in CA1, around six weeks post transduction (see Fig. 3). Optogenetic induction of SD via square wave light pulse (5000 ms, 488 nm, 5mW/mm$^2$, see fig. 3b). Movie played in real-time, the white trace shows simultaneous LFP recording in CA1. Note that the widefield imaging was blanked for the duration of the optogenetic stimulation from time 0-5 s.
