## Supplementary figures and images for "Region-specific spreading depolarization drives aberrant post-ictal behavior"

### Suppl_Fig01

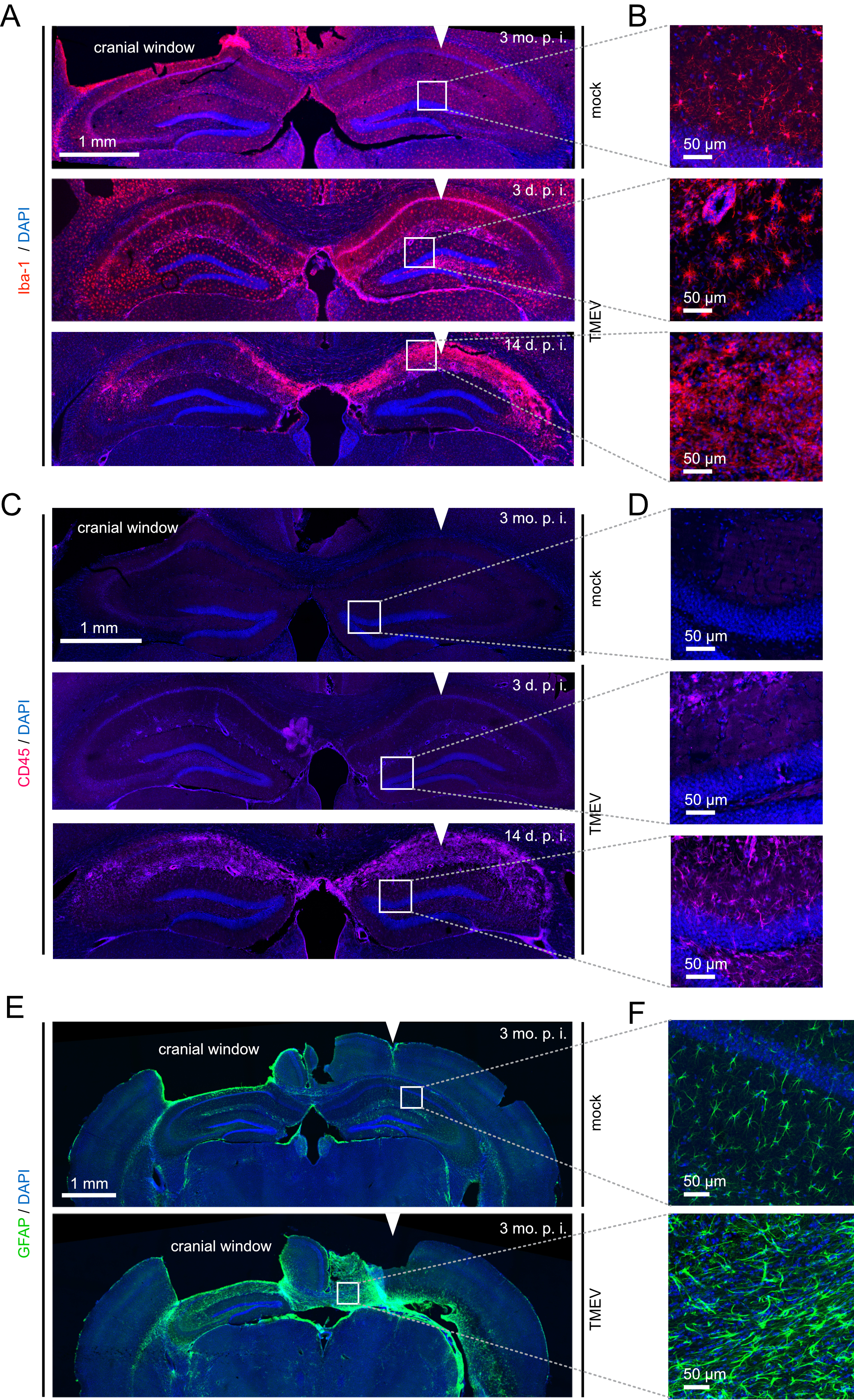

### Suppl_Fig02

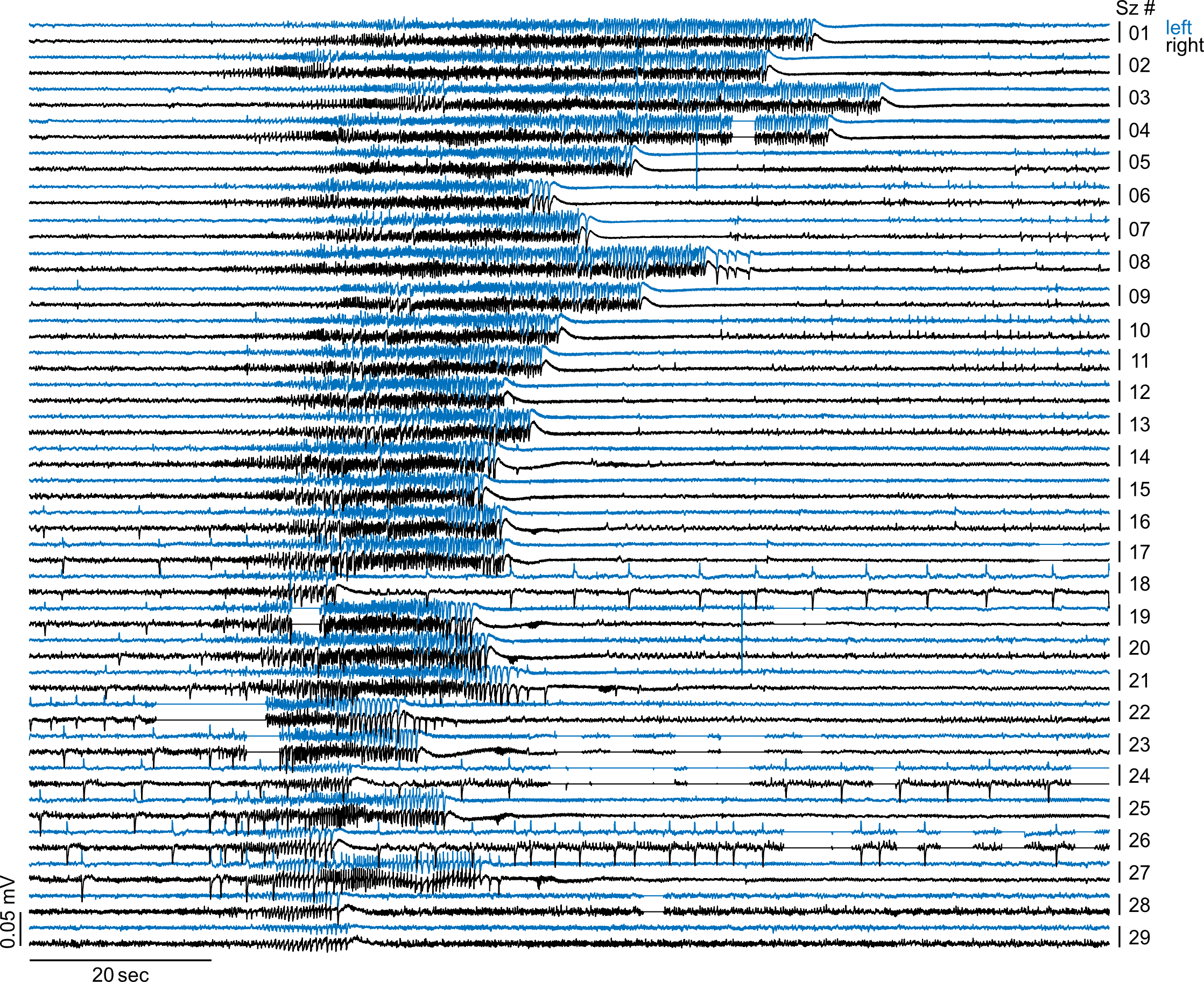

### Suppl_Fig03

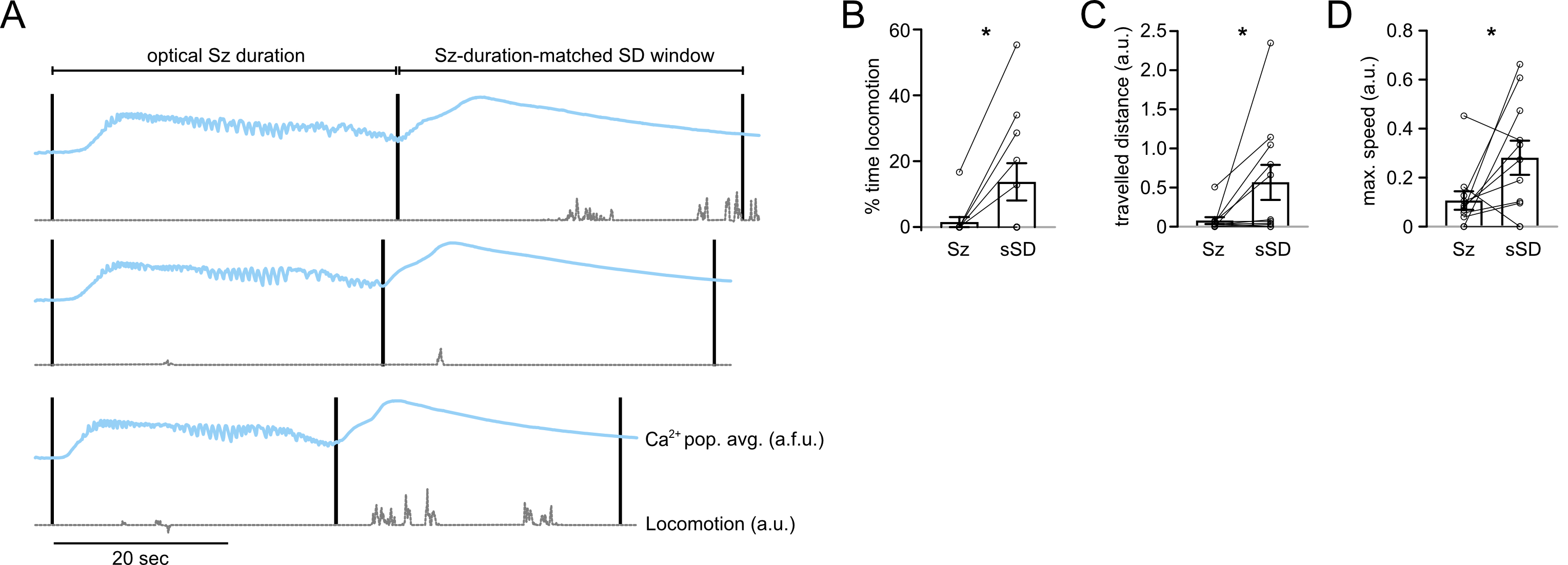

### Suppl_Fig04

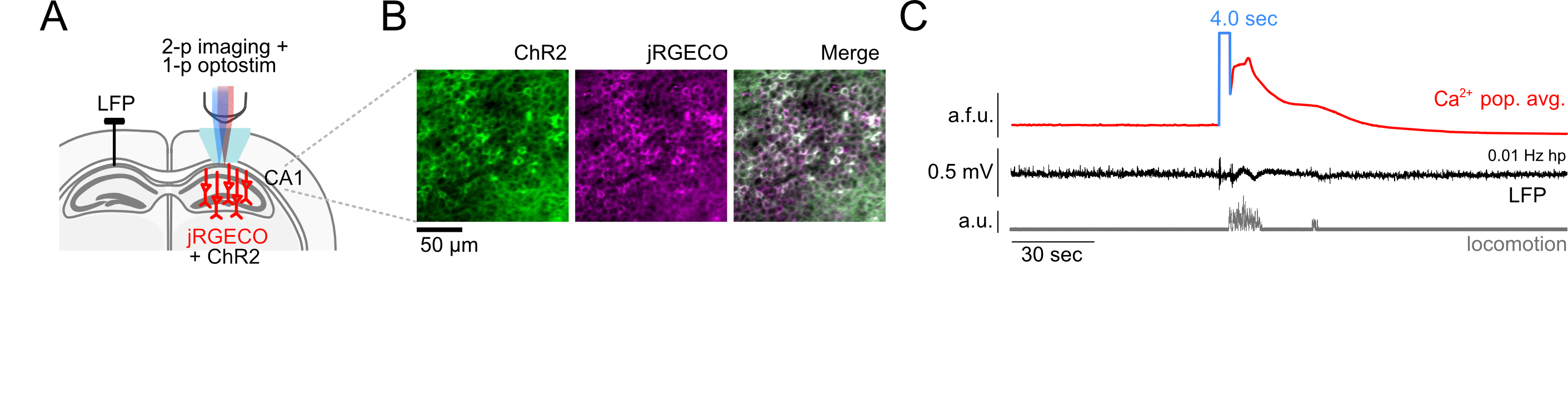

### Suppl_Fig05

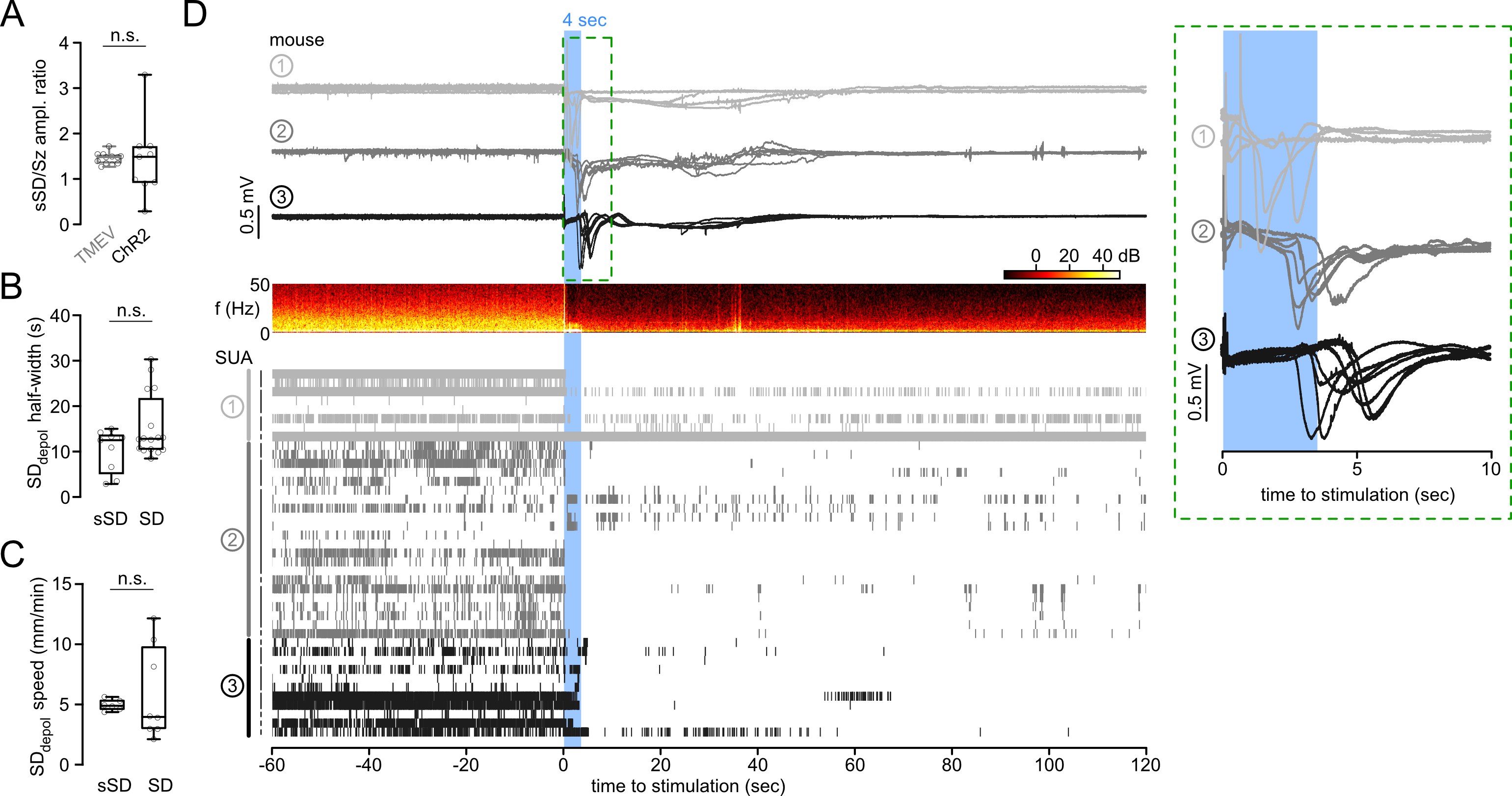

### Suppl_Fig06

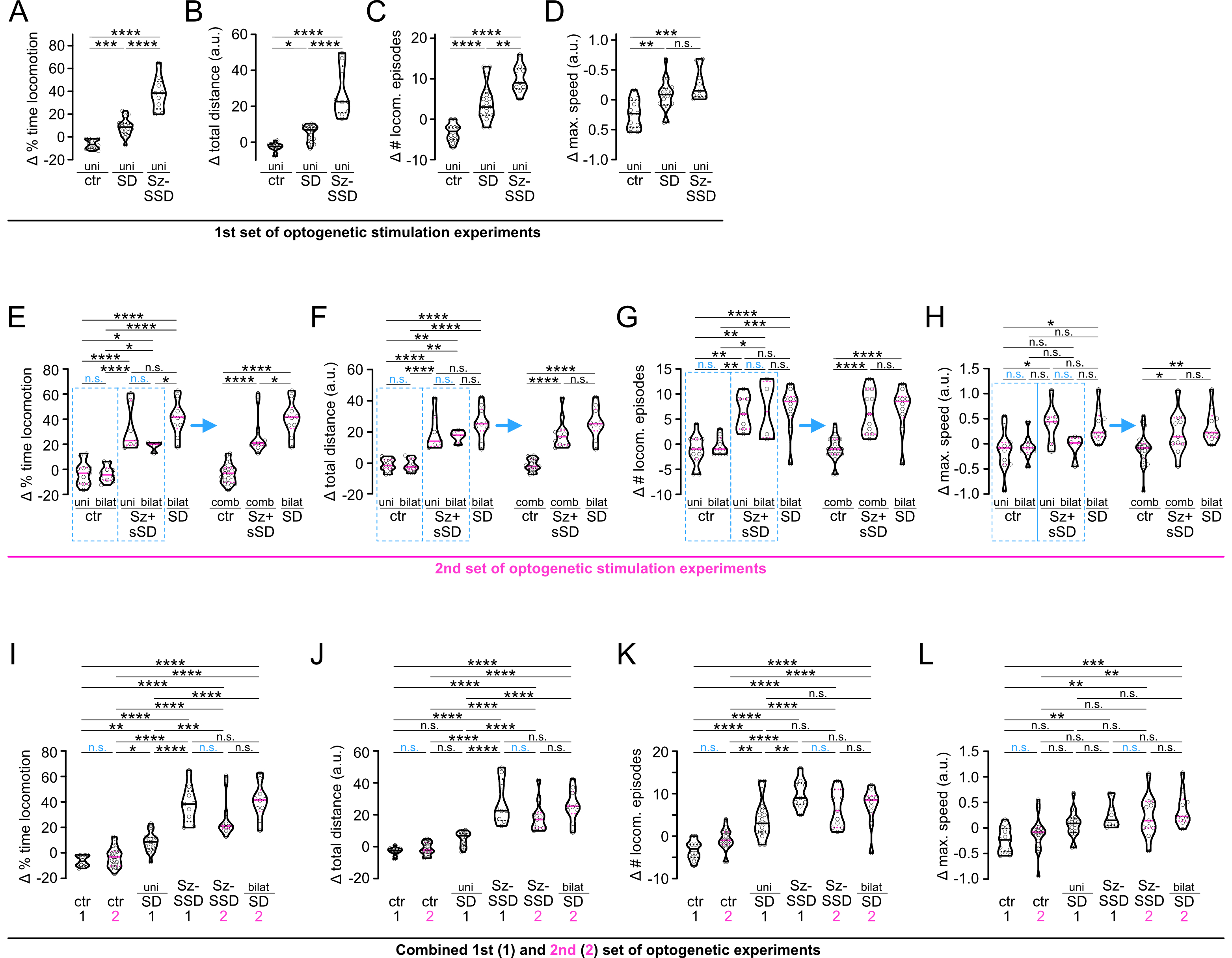

### Suppl_Fig07

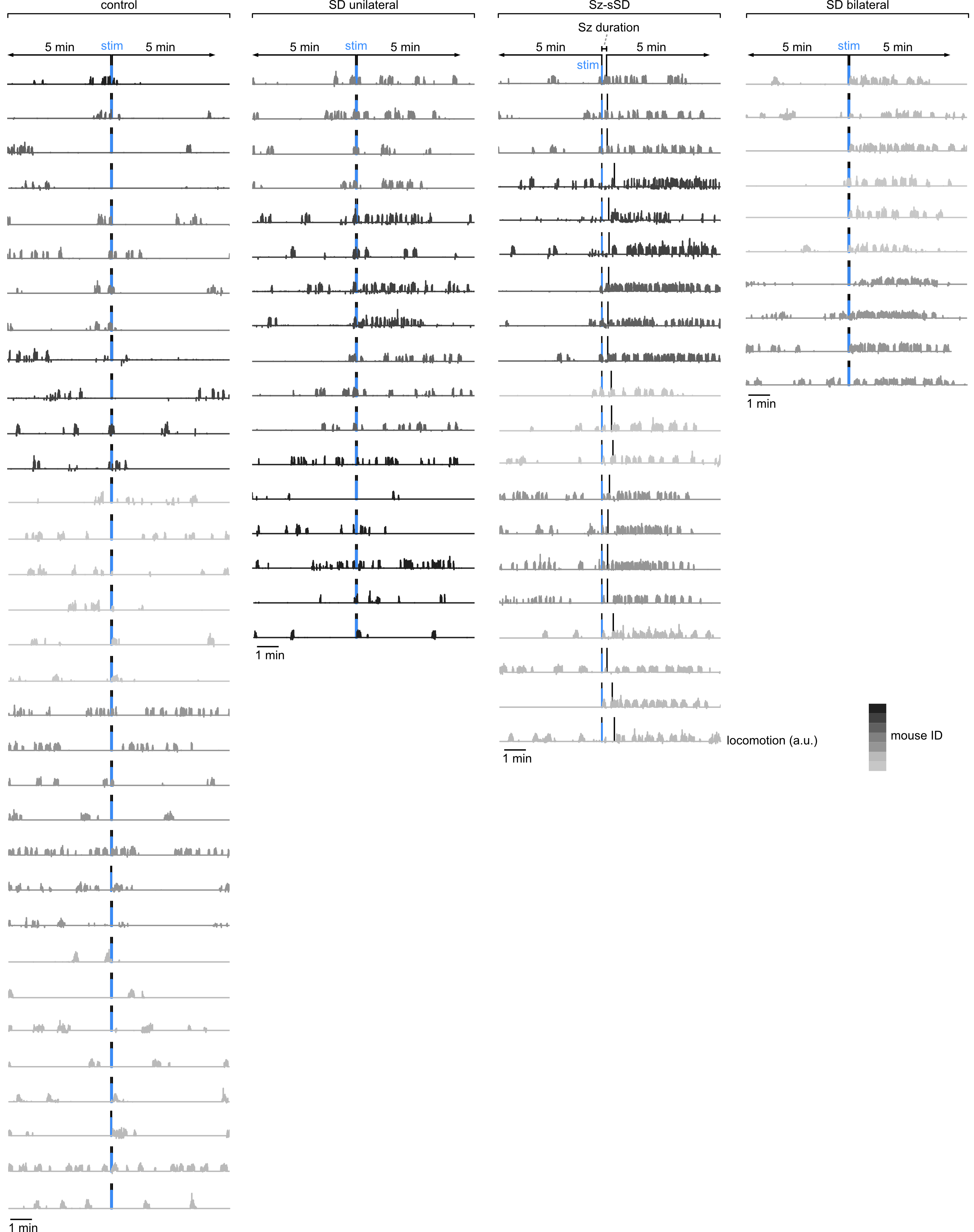

### Suppl_Fig08

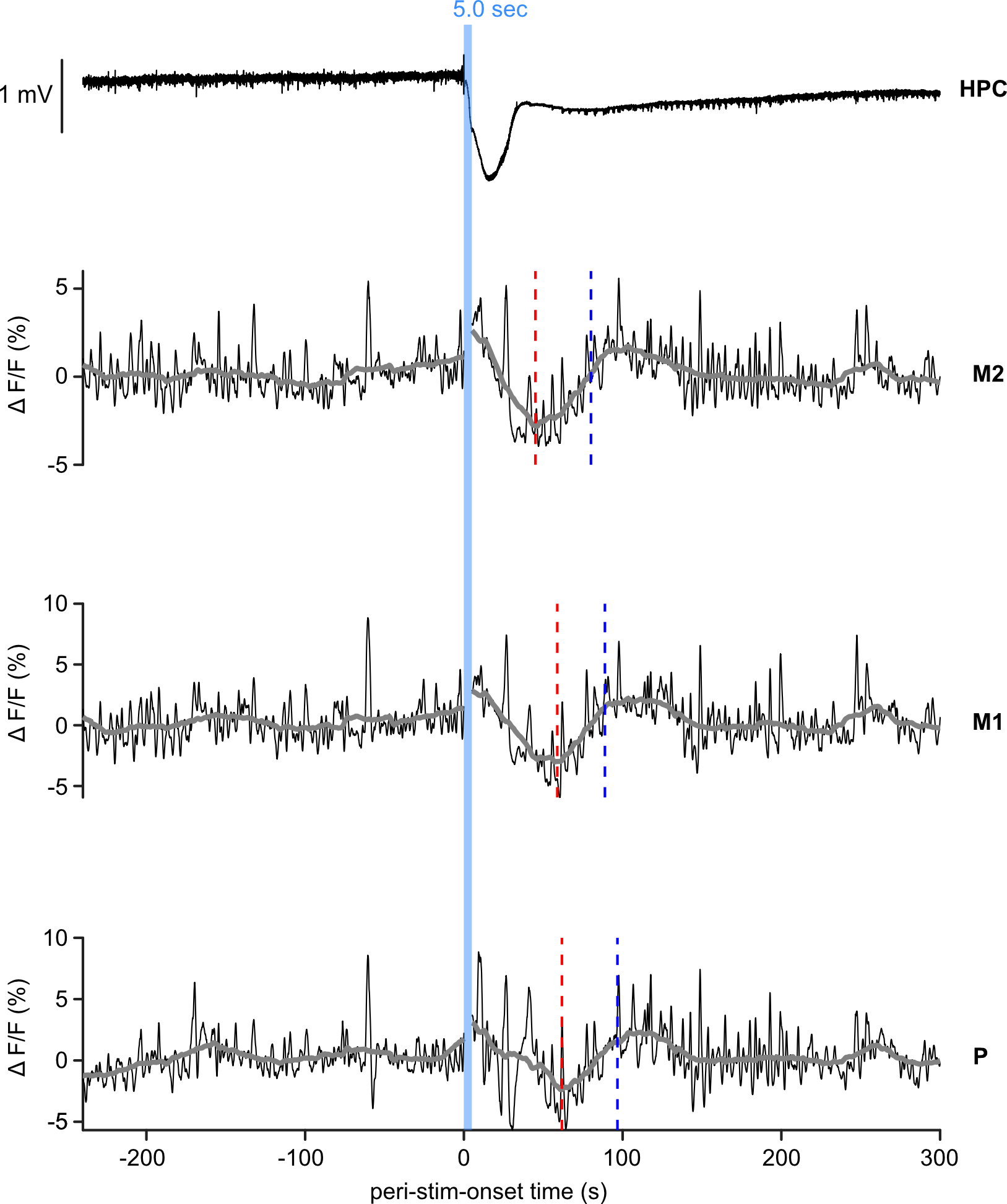

### Suppl_Fig09

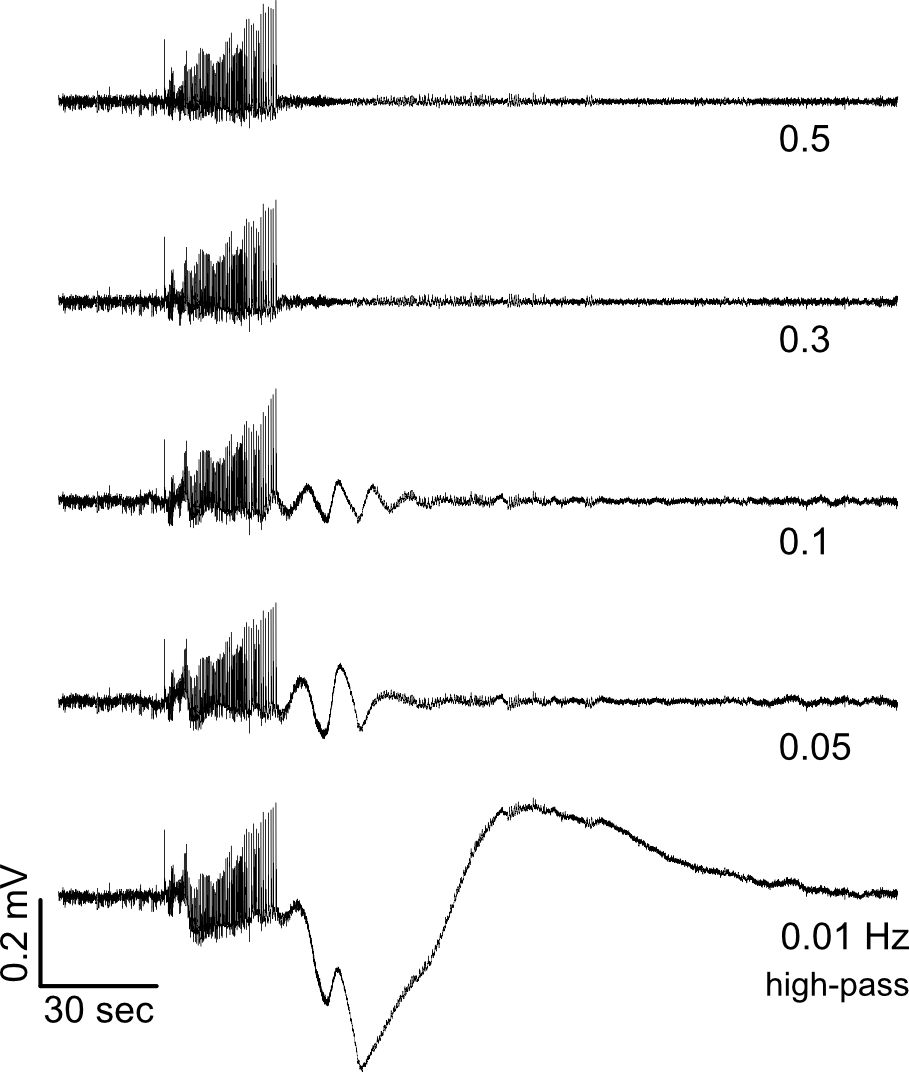

### Suppl_Fig10

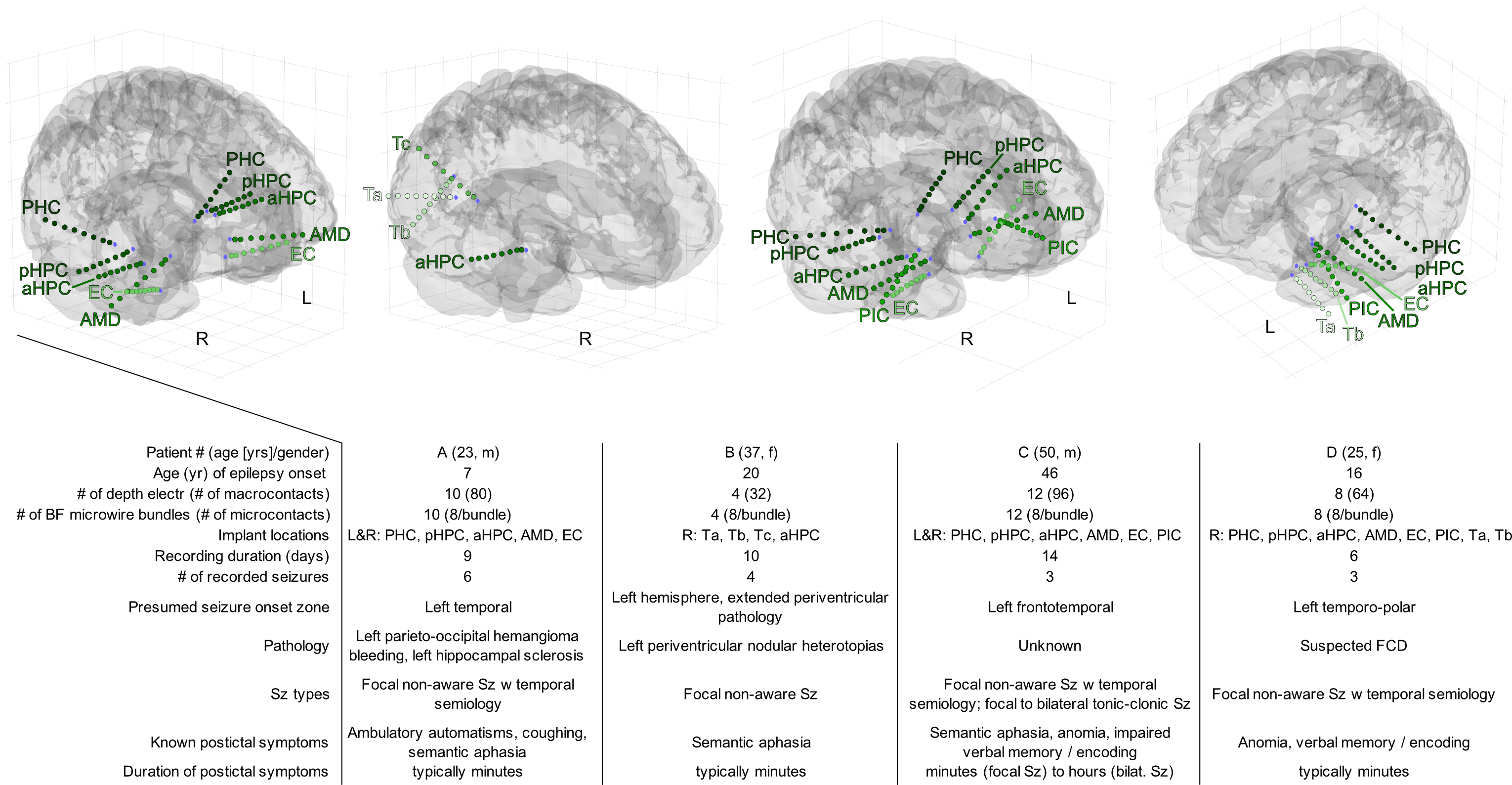

### Suppl_Table01

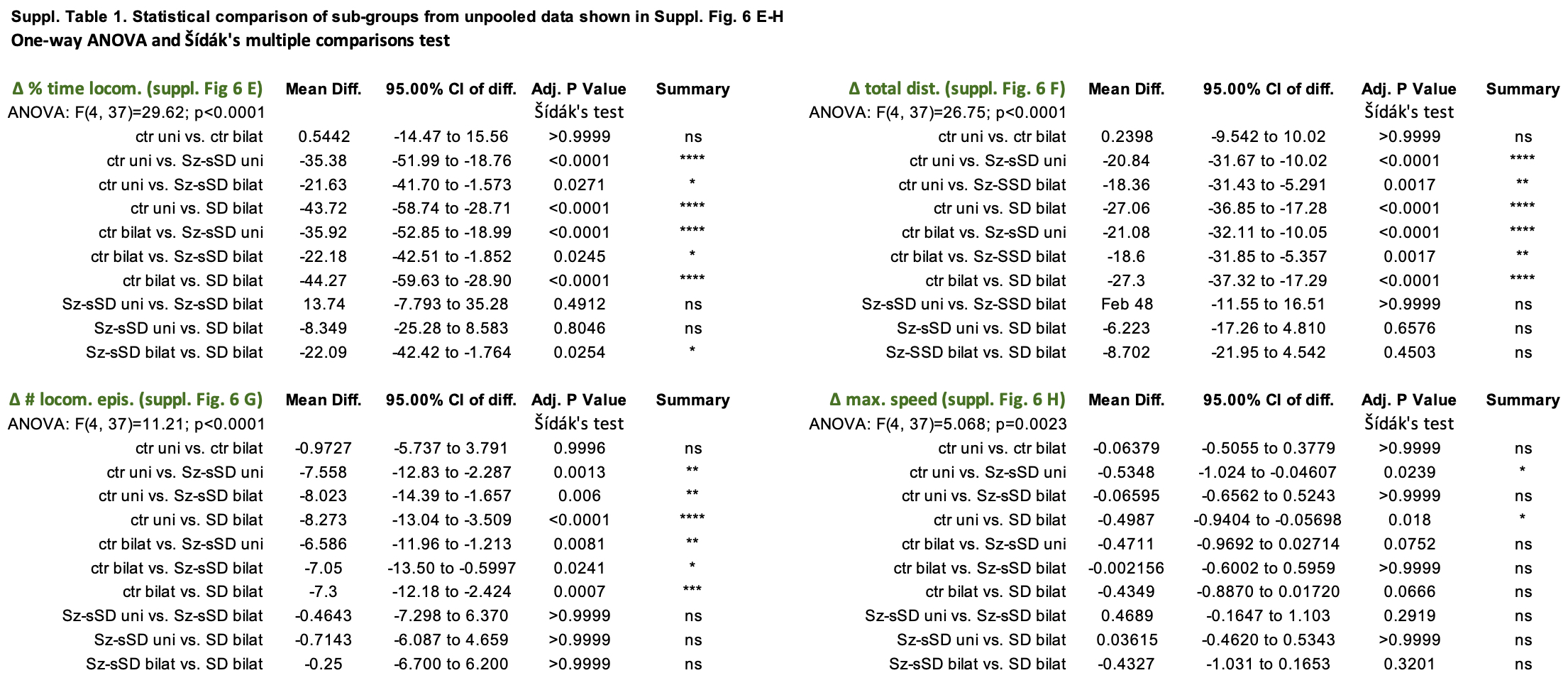

### Suppl_Table02

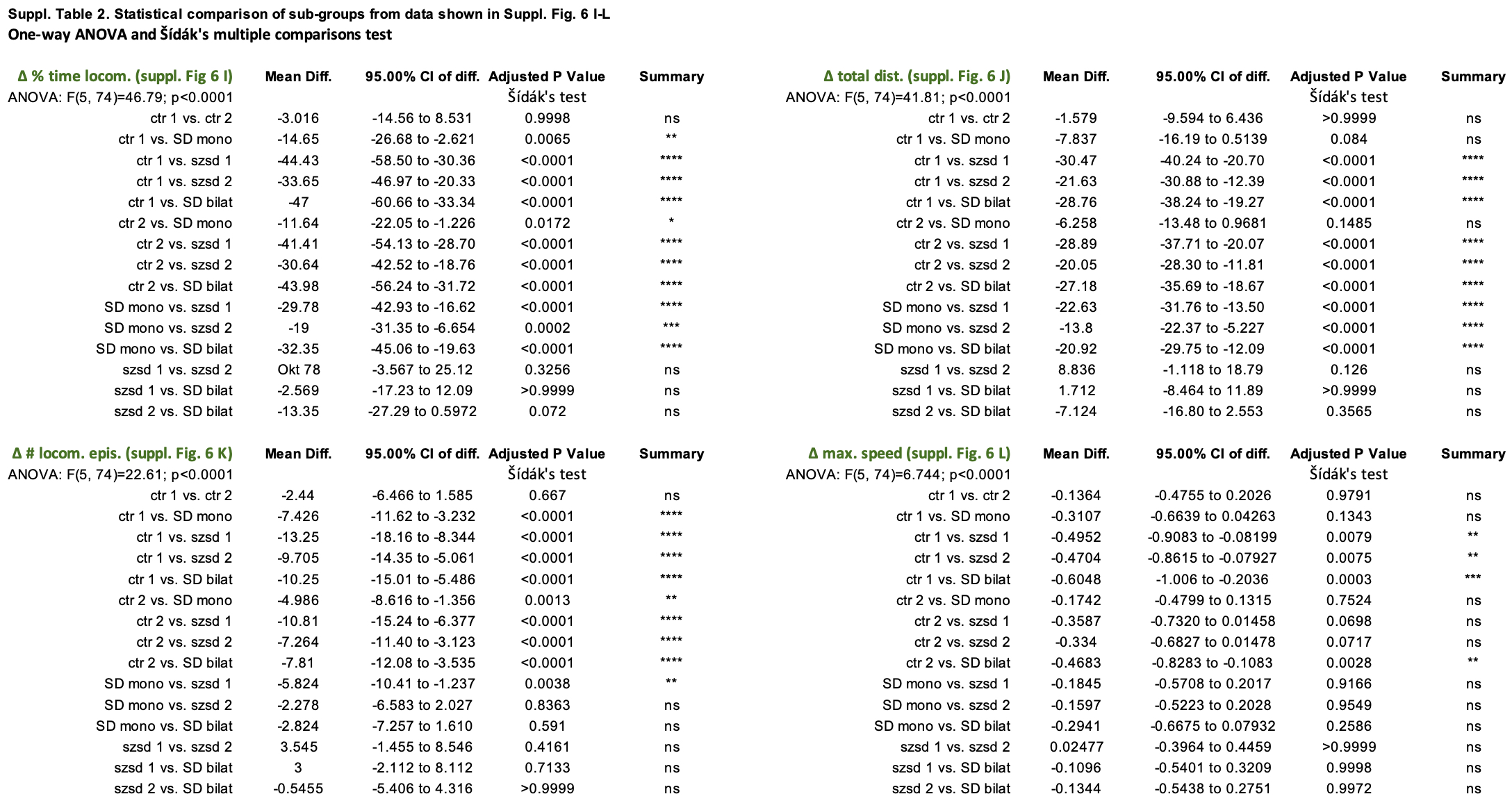
